# The gut microbiome stability of a butterflyfish is disrupted on severely degraded Caribbean coral reefs

**DOI:** 10.1101/2020.09.21.306712

**Authors:** Friederike Clever, Jade M. Sourisse, Richard F. Preziosi, Jonathan A. Eisen, E. Catalina Rodriguez Guerra, Jarrod J. Scott, Laetitia G.E. Wilkins, Andrew H. Altieri, W. Owen McMillan, Matthieu Leray

## Abstract

Environmental degradation has the potential to alter key mutualisms that underline the structure and function of ecological communities. While it is well recognized that the global loss of coral reefs alters fish communities, the effects of habitat degradation on microbial communities associated with fishes remain largely unknown despite their fundamental roles in host nutrition and immunity. Using a gradient of reef degradation, we show that the gut microbiome of a facultative, coral-feeding butterflyfish (*Chaetodon capistratus*) is significantly more variable among individuals at degraded reefs with very low live coral cover (~0%) than reefs with higher coral cover (~30%), mirroring a known pattern of microbial imbalance observed in immunodeficient humans and other stressed or diseased animals. We demonstrate that fish gut microbiomes on severely degraded reefs have a lower abundance of Endozoicomonas and a higher diversity of anaerobic fermentative bacteria, which suggests a broader and less coral dominated diet. The observed shifts in fish gut bacterial communities across the habitat gradient extend to a small set of potentially beneficial host associated bacteria (i.e., the core microbiome) suggesting essential fish-microbiome interactions are vulnerable to severe coral degradation.

## Introduction

Environmental degradation associated with the Anthropocene is threatening the persistence of mutualistic relationships that are key to the stability of ecological functioning^1^. The increasingly severe degradation of coral reefs from both local and climatic stressors has led to novel habitat states with conspicuously altered fish and invertebrate communities^2–5^, making them a model system for studying ecological responses to environmental change. A potentially pervasive but largely overlooked response to habitat degradation is the change to host-associated microbiomes - the communities of bacteria, archaea, fungi, unicellular eukaryotes, protozoa and viruses that live on internal and external surfaces of reef organisms. It has been suggested that coral microbiomes respond faster than their hosts to changing environmental conditions and can promote acclimatisation processes as well as genetic adaptation^6,7^. Microbial communities could play a key role in mediating a host’s resilience and ability to adapt to environmental change. However, it remains to be explored whether mutualisms between fish hosts and gut microbiomes can shift to alternative beneficial relationships to provide a mechanism of resilience to habitat change, or whether the mutualism breaks down and simply reflects a cascading effect of degradation at all levels of ecological organization. The importance of gut microbial communities in maintaining host health is well recognized in mammals and other vertebrates^8,9^, including a wealth of research into the importance of microbes in fish in aquaculture settings^10–13^. In coral reef fishes, recent studies have suggested that intestinal microbiomes influence key physiological functions associated with nutrient acquisition, metabolic homeostasis and immunity^14–17^. For example, gut bacteria provide many herbivorous fish hosts with the ability to digest complex algal polymers^14,15,18^. The gut microbiome is also a major actor in the innate immune responses to a wide variety of pathogenic microorganisms and other stressors in the surrounding environment^19,20^. Given the rapid physical, chemical and biotic changes affecting coral reefs, it is essential to gain a predictive understanding of how fish gut microbiome assemblages and metabolic functions respond to environmental variation to assess how the response of these mutualisms govern host health and resilience to habitat change.

Fish harbour microbiomes that are unique from the microbial communities in their surrounding environment^21,22^. The development of the gastrointestinal microbiome can start during hatching via acquisition of microorganisms from the egg’s chorion (i.e., the acellular protective envelope encasing the oocyte)^23^, and with 10,24–26 both water and the first food source entering the gastrointestinal tract^10,24–26^. Parental effects and host genotype likely mediate the early microbiome colonization process from egg and environmental sources^10,21,27^. As the gut microbiome diversifies throughout the development of the fish host, a relatively stable gut microbiome is typically established within the first months of the fish’s life and is influenced by a combination of host selection mechanisms and bacterial regulation of the fish host’s gene expression^12,19,28^. These resident (autochthonous) microbes, which are consistently found associated with the fish population across space and time and potentially provide critical functions for the host are referred to as the “core microbiome”^11,29,30^. In contrast, the numerous microbes occurring in the gastrointestinal tract after being ingested are transient (allochthonous) and may vary intraspecifically with developmental stage and potentially include opportunistic pathogens that may colonise in the case of impaired residential communities.

Because of their importance in maintaining host metabolic homeostasis, the degree of stability of the core microbiome across a range of environmental conditions is a key trait for predicting the resilience of the host population^31–33^. The stability of the core gut community may be altered if the host experiences severe physiological stress. It may switch to an alternative stable state (i.e., a novel but stable community), or communities may become more variable between individuals (i.e., communities are destabilized as stressors reduce the ability of hosts or their microbiome to regulate community composition)^34^.

The Chaetodontidae family (Butterflyfishes) is among the largest and most iconic families of coral reef-associated fishes^35^ and an ideal group for studying microbiome responses to habitat degradation. Chaetodontidae species range from extreme diet specialists to facultative corallivores and generalists capable of consuming different types of prey such as corals, algae, polychaetes or crustaceans^36–38^. Due to their intimate link to the reef benthos, specialized coral feeding species of Indo-Pacific butterflyfishes were shown to be highly sensitive to reductions in coral cover^39–41^. The foureye butterflyfish, *Chaetodon capistratus* (Linnaeus, 1758), is the only one of the four Western Atlantic *Chaetodon* species with a relatively high proportion of anthozoans in its diet (mainly hard and soft corals)^42–44^. Due to this relative specialization, we chose it as a model species to study links between reef habitats, hosts and the gut microbiome.

Here, we examined how differences in benthic habitat composition and coral coverage influence the variability and composition of the gut microbiota of *Chaetodon capistratus* across a tropical coastal lagoon in Bocas del Toro on the Caribbean coast of Panama. The Bay of Almirante encompasses an inner bay of protected reefs subjected to seasonal high temperatures and a watershed delivering nutrients from agriculture and sewage. In 2010, the bay faced an unprecedented hypoxic event, which led to massive coral bleaching and mortality on some reefs while others located near the bay’s mouth remained unaffected^45^. We capitalized on this gradient of habitat states across the bay of Almirante to test the resilience of fish gut microbiomes to environmental degradation. We hypothesized that fish residing on more degraded reefs have a more diverse microbiome as a result of alternative feeding behaviours and potentially increased stress^34^. On the other hand, we predicted that the core microbial community remains stable across the habitat gradient.

## Methods

### Study area

Bahia Almirante, located in the Bocas del Toro Archipelago on the Caribbean coast of Panama, is a coastal lagoonal system of approximately 450 km^2^ where numerous, relatively small and patchy fringing coral reefs occur^46^. Hydrographic and environmental conditions vary across the semi-enclosed bay but are generally characterized by limited water exchange with the open ocean^45^. Furthermore, areas of the bay are subjected to uncontrolled sewage and dredging due to increasing coastal development and agricultural runoffs from the adjacent mainland^47–50^. A total of nine discontiguous reefs distributed from the mouth towards the inner bay were selected for this study based on distinct hydro-geographical zones and disturbance history (Fig. 1A). Throughout the manuscript, we will refer to the three distinct reef zones as “outer bay”, “inner bay”, and “inner bay disturbed”. Outer bay reefs [Salt Creek (SCR), Cayo Corales (CCR) and Popa (PPR)] are located at the mouth of the bay. These reefs represent typical Caribbean reef communities featuring both massive and branching coral colonies with higher benthic cover and diversity as compared to the inner bay (Fig. 1B). Inner bay reefs [Almirante (ALR), Cayo Hermanas (SIS), and Cayo Roldan (ROL)] are largely coral and sponge dominated reefs of lower coral diversity than the outer bay reefs (Fig. 1C). Inner bay disturbed reefs [Punta Puebla (PBL), Punta STRI (PST) and Runway (RNW)] were heavily impacted by the 2010 hypoxic event (Altieri et al., 2017), which resulted in the current cover of largely dead coral comprised of formally prevalent *Agaricia* and *Porites* species (Fig. 1D).

**Figure 1.**
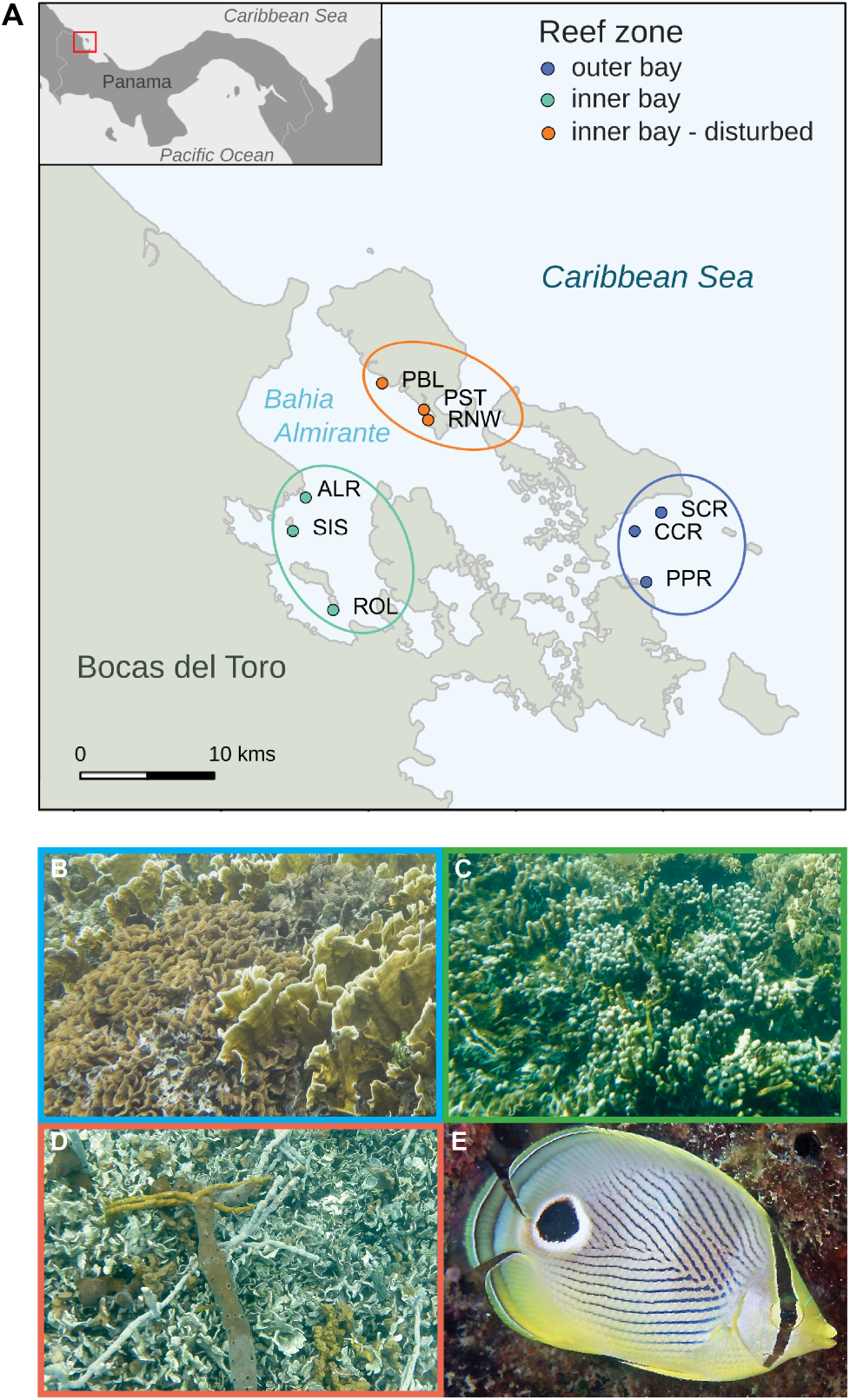
Study area and study species. (A) Map of the Bay of Almirante (Bocas del Toro, Panama) indicating the position of the nine reefs where samples were collected. (B) Inner bay reefs with intermediate levels of coral cover, (C) reefs located in the inner bay disturbed zone were highly impacted by a hypoxic event in 2010, (D) outer bay reefs with highest levels of live coral cover. (E) The study species foureye butterflyfish (*Chaetodon capistratus*).

### Benthic habitat and fish communities

Visual surveys of benthic cover and fish communities were conducted between May and June 2016. At each of the nine reefs, three 20 m transects were placed parallel to the shore at 2-4 m depth. Benthic community cover was estimated from 100 x 70 cm photographic quadrats placed every two meters resulting in a total of 10 quadrats per transect. Photos were analysed on the CoralNet platform^51^ using a stratified random sampling design (10 rows x 10 columns with 1 point per cell for a total of 100 points per image). Due to the difficulty involved with photo-based taxonomic identification, analyses were conducted at the level of broad functional groups. Mean cover of each benthic category was calculated per reef. Fish communities were characterized by one trained surveyor who recorded the identity and abundance of all reef fishes encountered along each 20 m belt (2.5 m on each side of the transect line) while swimming slowly using scuba (except at CCR).

### Sample collection

The foureye butterflyfish, *Chaetodon capistratus,* is a common member of Caribbean coral reef fish communities (IUCN classified as least concern)^52^ with a distribution that extends across the subtropical Western Atlantic^53–55^ (Fig. 1E). The following protocol of fish capture and euthanization was approved by the Smithsonian Tropical Research Institute’s Institutional Animal Care and Use Committee (IACUC). An average of 11 individual adult fish were collected at each of the nine reefs (min = 7; max = 16; total = 102) by spearfishing in February and March 2018 (Table S1). Captured fish were immediately brought to the boat, anesthetized with clove oil and placed on ice in an individual and labelled sterile Whirl-Pak bag. Upon return to the research station, fish were weighed (g wet weight), and Standard Length (mm SL) as well as Total Length (mm TL) were measured with a digital caliper. The intestinal tract of each fish was removed under a laminar flow hood using tools decontaminated with 10% sodium hypochlorite, preserved in 96% ethanol in individual 15 ml or 5 ml centrifuge tubes and stored at −20°C until DNA extraction. To assess microbial communities present in the fish’s environment, we also obtained samples of potential prey taxa and seawater. At each of nine reefs, a total of four liters of seawater was collected immediately above the reef substratum using sterile Whirl-Pak bags and filtered through a 0.22 μm nitrocellulose membrane (Millipore). Small pieces of hard coral (*Siderastrea siderea, Porites furcata, Agaricia tenuifolia*), soft coral (*A. bipinnata, Plexauridae* sp.), sponges (*Amphimedon compressa, Niphatidae* sp., *Chondrilla caribensis, Poecilosclerida* sp., *Chalinidae* sp., *Xestospongia* sp.), macroalgae (*Amphiroa* sp.), turf, and zoantharia (*Zoanthus pulchellus, Palythoa caribaoerum*) were collected and kept in sterile Whirl-Pak bags on ice on the boat. At the field station, samples were individually placed in 50 ml or 15 ml centrifuge tubes with 96% ethanol and stored at −20 °C until DNA extraction.

### DNA analysis

The gastrointestinal tract of each fish was opened longitudinally to isolate the digesta and the mucosa tissue by lightly scraping the intestinal epithelium. Between 0.05 and 0.25 g of both tissue types combined was used for DNA extraction using the Qiagen Powersoil DNA isolation kit following the manufacturers instructions with minor modifications to increase yield (see Supplementary text). Tissues of all potential prey organisms (invertebrates and macroalgae) were homogenized per sample. Additionally, infaunal communities (small worms) were isolated from two sponges, *Amphimedon compressa* and *Dysidea sp*. and the tissue homogenized for each sponge separately. DNA was extracted (0.25g per sample) following protocols described previously. Seawater DNA was isolated from nitrocellulose membranes filters using the Qiagen Powersoil Kit following a modified protocol described previously^56^.

A dual Polymerase Chain Reaction (PCR) approach was used to amplify the V4 hypervariable region (primers 515F^57^ and 806R^58^) of the 16S ribosomal rRNA gene of each sample and the product of all samples was sequenced by combining into a single Illumina MiSeq sequencing run. Our protocol followed the 16S Illumina Amplicon Protocol of the Earth Microbiome Project^59^ using locus-specific primers to which Illumina “overhang” sequences were appended. These overhang sequences served as a template to add dual index Illumina sequencing adaptors in a second PCR reaction (see supplementary text for detailed PCR protocols). The final product was sequenced on the Illumina MiSeq sequencer (reagent kit version 2, 500 cycles) at the Smithsonian Tropical Research Institute with 20% PhiX. The absence of contaminants was confirmed with negative DNA extractions and negative PCR amplifications (see supplementary text for detailed DNA extraction and PCR protocols).

### Analysis of sequence data

Illumina adapter and primer sequences were removed from forward and reverse reads using “cutadapt”^60^ with a maximum error rate of 0.12 (-e 0.12). Remaining reads were filtered and trimmed based on their quality profiles and potential chimeras removed using DADA2 1.12.1^61^ in R environment version 3.6.1^62^. Sequences were discarded if they had more than two expected errors (maxEE = 2), or at least one ambiguous nucleotide (maxN = 0) or at least one base with a high probability of erroneous assignment (truncQ = 2). Forward and reverse reads were trimmed to 220 bp and 180 bp respectively to remove lower quality bases while maintaining sufficient overlap between paired end reads. Sequences were kept when both the forward and reverse reads of a pair passed the filter. Quality filtered reads were dereplicated and Amplicon Sequence Variants (ASVs) inferred. Paired-end reads were merged and pairs of reads that did not match exactly were discarded. Taxonomy was assigned to each ASV using a DADA2 implementation of the naive Bayesian RDP classifier^63^ against the Silva reference database version 132^64^. ASVs identified as chloroplast, mitochondria, eukaryota, or that remained unidentified (i.e, “NA”) at the kingdom level were removed from the dataset. Sequences of each ASV were aligned using the DECIPHER R package version 2.0^65^. The phangorn R package version 2.5.5^66^ was then used to construct a maximum likelihood phylogenetic tree (GTR+G+I model) using a neighbor-joining tree as a starting point. Fourteen samples containing few sequences were removed from the dataset (Fig. S1). The remaining samples were rarefied to even sequencing depth (n = 10,369 sequences) for downstream analysis. Our approach followed the recommendation for normalization of sequencing data^67^. Statistical analysis was conducted using phyloseq version 1.28.0 in R^68^.

### Delineation of the core gut microbiome

To identify the persistent and potentially beneficial bacteria associated with the fish gut (i.e., the “core gut microbiome”^29,69^), we employed a statistical approach taking into account both relative abundance and relative frequency of occurrence of ASVs as opposed to the common procedure of using an arbitrary minimum frequency threshold 69 70 71 based on presence-absence data only^69^. Indicator species analysis^70^ (labdsv package)^71^ was used to identify which ASVs were relatively more abundant and predominantly found in fish guts and not in their surrounding environment. We calculated an Indicator Value (IndVal) Index between each ASV and two groups of samples: (1) all fish gut samples, and (2) all seawater and sessile invertebrate samples upon which the fish potentially feeds (Fig. 2). The statistical significance of the association between ASVs and groups of samples was tested using 1000 permutations. ASVs were considered indicators of fish guts (i.e., components of the core) if *P*-value ≤ 0.001.

**Figure 2.**
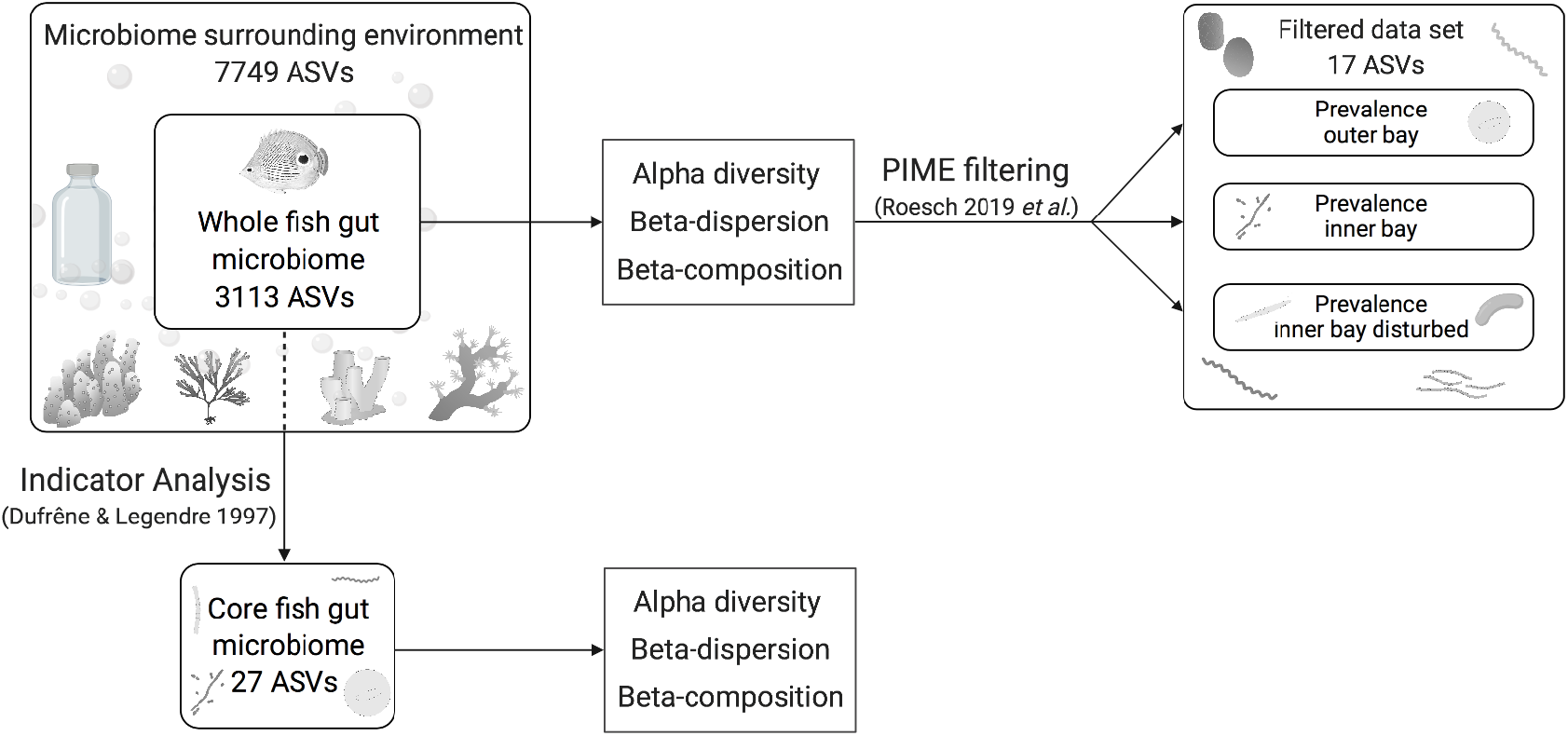
Microbial community analysis workflow illustrating how we subsetted the whole fish gut microbiome dataset to delineate the core microbiome and microbial zone communities, respectively. To identify the core microbiome, we used indicator analysis between the whole fish gut microbiome and the environmental sample fraction consisting of samples of potential fish prey taxa and the surrounding seawater. Diversity analysis was done for the whole and core fish gut microbiome, respectively. The whole fish gut microbiome was filtered for prevalence with a machine learning-based algorithm (PIME)^87^ to detect community differences among zones that reflect fish microbiome responses to the habitat gradient.

Sequences of ASVs identified as part of the core microbiome were compared to the non-redundant nucleotide (nr/nt) collection database of the National Centre for Biotechnology Information (NCBI) using the Basic Local Alignment Search Tool for nucleotides (BLASTn)^72^. We extracted metadata associated with all sequences that matched each query at 100% similarity or the first five top hits to identify where each core ASVs and close relatives were previously found.

### Diversity analyses

The workflow of our microbial community analysis is visualized in a diagram (Fig. 2). To account for presence of rare sequence variants caused by sequencing errors or other technical artifacts^73^, we used Hill numbers^74^ following the approach recommended by Alberdi & Gilbert (2019)^75^ for sequence data to compare alpha diversity between groups of samples. Hill numbers allow scaling the weight put on rare versus abundant sequence variants while providing intuitive comparisons of diversity levels using “effective number of ASVs” as a measuring unit^74–76^. This approach allowed for balancing the over representation of rare ASVs that might be inflated due to sequencing errors^77^. We calculated three metrics that put more or less weight on common species: (1) observed richness, (2) Shannon exponential that weighs ASVs by their frequency, and (3) Simpson multiplicative inverse that overweighs abundant ASVs. Alpha diversity was calculated and visualized using boxplots for the whole and core fish microbiome. Because Shapiro-Wilk tests indicated that the data were not normally distributed, non-parametric Kruskal-Wallis tests were used to compare alpha diversity among reefs (N=9) and the three reef zones (outer bay, inner bay, inner bay – disturbed) with post-hoc Dunn tests.

To test the hypothesis that fish gut microbiome are more variable between individuals at disturbed sites, we calculated non-parametric Permutational Analysis of Multivariate Dispersion (PERMDISP2) (betadisper function, vegan package implemented in phyloseq^68,78^). PERMDISP2 is a measure of the homogeneity of variance among groups and compares the average distance to group the centroid between each predefined group of samples in multidimensional space. We used a range of phylogenetic and non-phylogenetic dissimilarity metrics that differentially weigh the relative abundance of ASVs to identify the effect of abundant ASVs^79^ [Phylogenetic: Unifrac, Generalized Unifrac (package GUniFrac)^80^ and Weighted Unifrac^81^; non-phylogenetic: Jaccard^82^, modified Gower with log base 10^79^ and Bray Curtis^83^]. *P*-values were obtained by permuting model residuals of an ANOVA (Analysis of Variance) null-model 1000 times (betadisper function, vegan package implemented in phyloseq^68,78^). Principal Coordinates Analysis (PCoA) plots were generated for each distance measure respectively to visually explore patterns of variance dispersion across the three reef zones.

Differences in microbial composition were tested using Permutational Multivariate Analysis of Variance (PERMANOVA) with the Adonis function in vegan^84^ computed with 10,000 permutations. Comparisons were made (1) between fish gut microbiomes of the three reef zones (‘zone model’), (2) between fish gut microbiomes of outer bay reefs versus inner bay reefs (‘position model’) and (3) between fish gut microbiomes of inner bay reefs and inner bay disturbed reefs which differed in coral cover (‘cover model’). Permanova is robust to the effect of heterogeneity of multivariate dispersions in balanced or near balanced designs as in our study^85^. Pairwise Adonis with Bonferroni corrected *P*-values was computed using the pairwise Adonis function in R^86^.

Finally, we used the Prevalence Interval for Microbiome Evaluation (PIME) R package^87^ to identify sets of ASVs that are predominantly found (more frequent) in fish guts of each zone of the Bay of Almirante (outer bay, inner bay, inner bay – disturbed). PIME uses a supervised machine learning Random Forest algorithm^88^ to reduce within-group variability by excluding low frequency sequences potentially confounding community comparisons of microbiome data^87^. PIME identifies the best model to predict community differences between groups by defining a prevalence threshold that retains as many ASVs as possible in the resulting filtered communities (i.e., the random forest classifications), while minimizing prediction error (out of bag error, OBB). To do so, the algorithm uses bootstrap aggregating (100 iterations) of each sample group at each filtering step (prevalence interval) by 5% increments. Random Forest calculates a global prediction from a multitude of decision trees based on the bootstrap aggregations and estimates the out of bag error rate (OBB) from omitted subsamples during aggregating^88^. Validation was done by randomizing the original dataset (100 permutations) and subsequently estimating Random Forest error to determine if group differences in the filtered dataset were due to chance (pime.error.prediction function, PIME package)^87^. A second function (pime.oob.replicate, PIME package)^87^ repeated the Random Forest analysis using the filtered dataset for each prevalence interval without randomizing group identity. In a preliminary step, we assessed whether the OOB error for our unfiltered data was >0.1, which indicated that de-noising with PIME would improve model accuracy.

## Results

### Benthic habitat and fish communities

Reefs located in the three zones classified *a priori* as outer bay, inner bay and inner bay disturbed, differed both in terms of their benthic composition (PCoA; Fig. 3A) and level of live coral cover (Fig. 3B). Live coral cover (mean cover per site: SCR 37.1%, PPR 33%, CCR 29.3%; Fig. 3B) and coral diversity (Shannon diversity; Fig. S2) were highest on reefs of the outer bay. Both stony coral species (i.e. *Acropora cervicornis, Agaricia tenuifolia*) and fire corals (i.e. *Millepora alcicornis, Millepora complanata*) dominated at outer bay reefs. At the inner bay zone, reefs displayed an intermediate level of live coral cover (mean cover per transect: ALR 21.2%, SIS 13.3%, ROL 9.4%; Fig. 3B), largely dominated by the lettuce coral *Agaricia tenuifolia*. Sponges represented more than a quarter of the benthic cover at these reefs (mean sponge cover per transect: ALR 23%, SIS 18.5%, ROL 34.2%; Table S3). Live coral cover was lowest at the inner bay disturbed zone (mean cover per transect: RNW 0.8%, PST 0.3%, PBL 0%; Fig. 3B) where dead coral skeleton was prevalent (mean cover per transect: RNW 45.3%, PST 21.4%, PBL 53.6%) together with sponges (mean cover per transect: RNW 27.3%, PST 21.3% and PBL 21.9%; Table S3). Principal Coordinates Analysis (PCoA; Bray Curtis dissimilarity) indicated distinct fish communities at the inner bay disturbed zone. In contrast, fish communities at the inner and outer bay zone appeared more similar (Fig. S3A). Our focal species *Chaetodon capistratus* was present at all surveyed reefs in similar abundance levels (1 - 5 individuals per 100 m^2^ transect) apart from Cayo Hermanas (SIS, inner bay zone) where up to 25 individuals were recorded in one of the transects (Fig. S3B).

**Figure 3.**
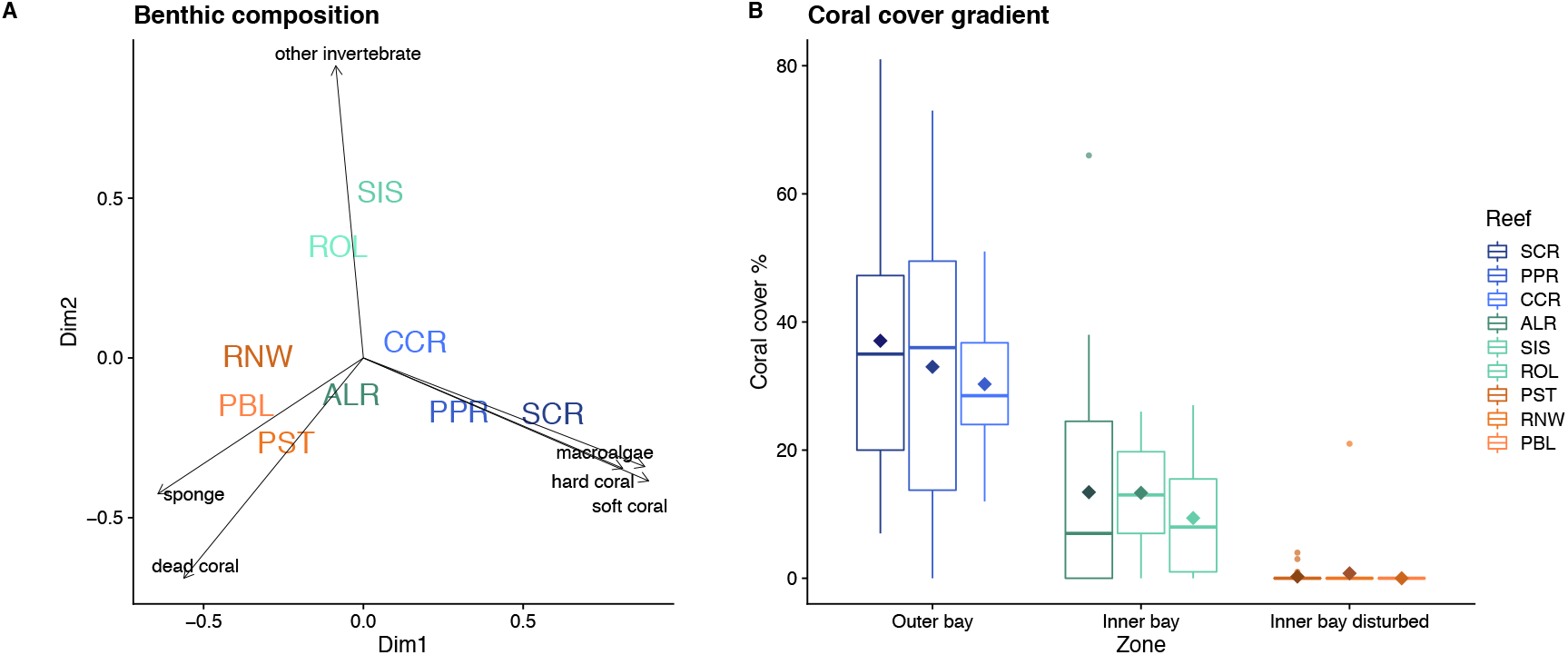
Composition and percent coral cover of benthic communities across nine reefs and three reef zones illustrating a habitat gradient: (A) PCoA representing dissimilarities in benthic community composition based on Bray-Curtis. Reefs are colour coded by reef zone, substrate groups are depicted in black; (B) percent live coral cover across reef zones from high coral cover at the outer bay to very low cover at disturbed reefs at the inner bay. Diamond shapes depict means.

### Composition of the whole gut microbiome

A total of 5,844,821 high quality reads were retained for subsequent analyses. The number of reads per sample ranged from 10,369 to 79,466, with a mean ± SD of 41,307 ± 10,990 reads. 10,711 different ASVs were identified in the total dataset. The number of ASVs per sample ranged from 13 to 1,281, with a mean ± SD of 179 ± 210 ASVs. This data set primarily comprised ASVs belonging to 15 bacterial phyla (abundance > 5%; Fig. S4A). As predicted, *C. capistratus’* gut microbiome composition was distinct from the microbiome in seawater and the microbiome of potential prey items (sessile invertebrates) (Fig. S4A and S4B). *Chaetodon capistratus’* overall gut microbiome was dominated by Proteobacteria (68.6%) followed by Firmicutes (16.1%), Spirochaetes (9.27%), Cyanobacteria (3.98%) (Fig. S4A). Bacteria in the phylum Proteobacteria were dominated by a single genus (*Endozoicomonas*) in the gut of *C. capistratus* (93.9%) (Fig. S4B). Firmicutes was abundant in fish guts (16.1% of fish gut bacteria) but representatives of this phylum were nearly absent from potential prey (0.005%, 0.06%, 0.47%, 0.26% and 0.07% in algae, hard corals, soft corals, sponges and zoanthids, respectively) and seawater (and 0.02%) (Fig. S4A and S4B).

### Composition of the core gut microbiome

Indicator Analysis identified 27 ASVs in eight families (i.e., Endozoicomonadaceae, Brevinemataceae, Ruminococcaceae, Lachnospiraceae, Vibrionaceae, Peptostreptococcaceae, Clostridiaceae, Thermaceae) as part of the ‘core’ microbiome associated with the fish intestinal tract (IndVal; *P* 0.001) (Fig. S5, Table S5A). The genus *Endozoicomonas* (phylum Proteobacteria, class Gammaproteobacteria), described as a symbiont of marine invertebrates (Naeve 2017), comprised 71.3% of the ASVs in the core followed by genus *Brevinema* (phylum Spirochaetes, class Spirochaetia) (10.7%) and anaerobic fermentative bacteria in the families Ruminococcaceae (9.7%), Lachnospiraceae (5.6%), and Clostridiaceae (1.7%) (phylum Firmicutes, class Clostridia).

Blastn searches against nr/nt NCBI database revealed that ASVs identified as part of the core gut microbiome were previously found in scleractinian and soft coral tissue (*Endozoicomonas* ASV1, ASV3, ASV5, ASV6, ASV11, ASV17) at our study area and in Curacao (ASV1, ASV3, ASV5, ASV17, ASV68) among other locations (ASV1, ASV5, ASV7, ASV11, ASV68) (Table 1). Some *Endozoicomonas* ASVs were closely related to sequences identified previously in sponges, clams, ascidians, tunicates, and coral mucus (ASV7, ASV59, ASV68, ASV 163) as well as the intestinal tract of a coral reef fish species (*Pomacanthus sexstriatus*) (ASV5). Sequences assigned to Ruminococcaceae closely resembled bacteria reported from herbivorous marine fishes (*Kyphosus sydneyanus, Naso tonganus, Acanthurus nigrofuscus, Siganus canaliculatus*) (ASV9, ASV14, ASV15, ASV25, ASV39), the omnivorous coral reef fish *Pomacanthus sexstriatus* (ASV25) and a freshwater fish (ASV18). An Epulopiscium ASV matched with 100% identity to a sequence detected in the guts of two coral reef fishes, the omnivorous *Naso tonganus* and the carnivorous *Lutjanus bohar* (ASV27) and to sequences found in the coral *Orbicella faveolata* (ASV27). Other Lachnospiraceae bacteria found in this study resembled sequences known from cattle rumen, hot springs, farm waste, human and other animal feces (ASV10, ASV 24). Within Ruminococcaceae in Firmicutes, ASVs assigned to genus *Flavonifractor* closely resembled bacteria reported from the hind gut of the temperate herbivorous marine fish *Kyphosus sydneyanus* in New Zealand (ASV9). *Brevinema* sequences similar to ours have been previously isolated from the gut of the coral reef fish *Naso tonganus* as well as freshwater and intertidal fish intestinal tracts (ASV2). Retrieved Vibrionaceae (genus *Vibrio*) were similar to sequences found in a coral reef fish gut of *Zebrasoma desjardinii* (ASV95). An *Romboutsia* ASV (family Peptostreptococcaceae), a recently described genus of anaerobic, fermentative bacteria associated with the intestinal tract of animals including humans^89–91^ but which also occur in mangrove sediments^92^ matched 100% a sequence found in tissue of the sea fan *Gorgonia ventalina* at our study site Bocas del Toro (ASV 30) (Table 1).

**Table 1.**
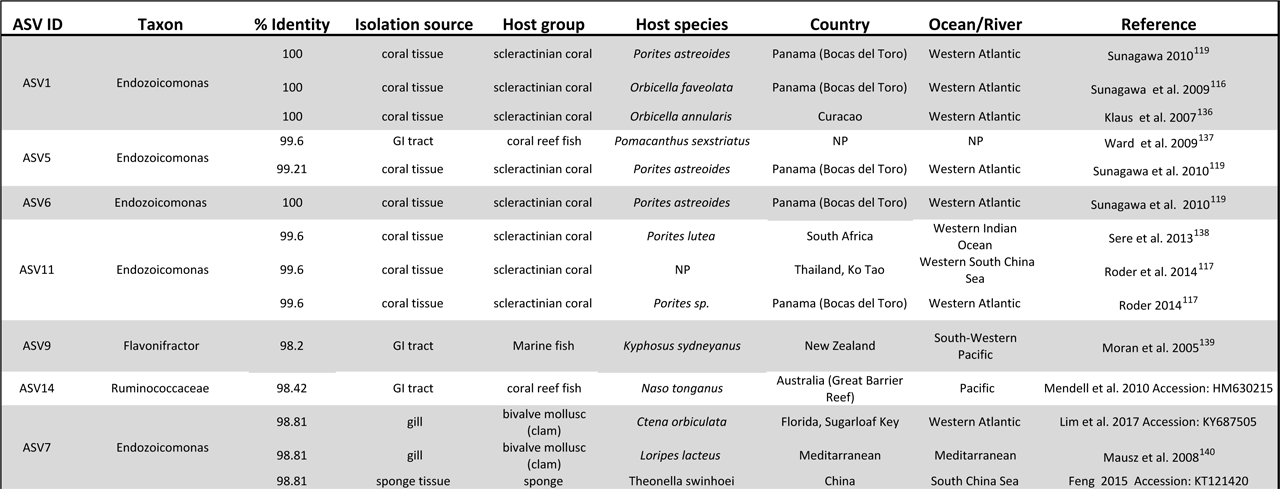
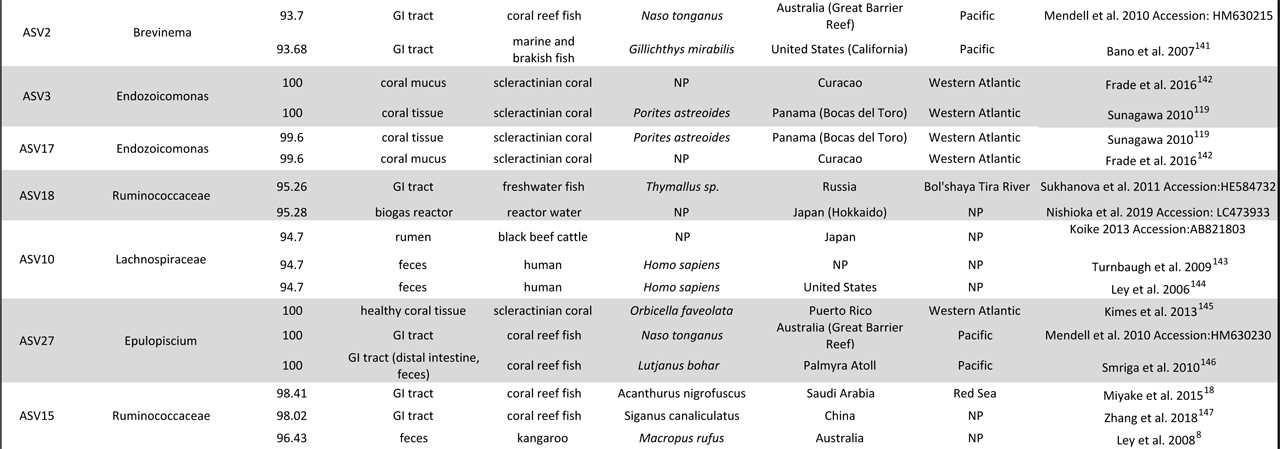
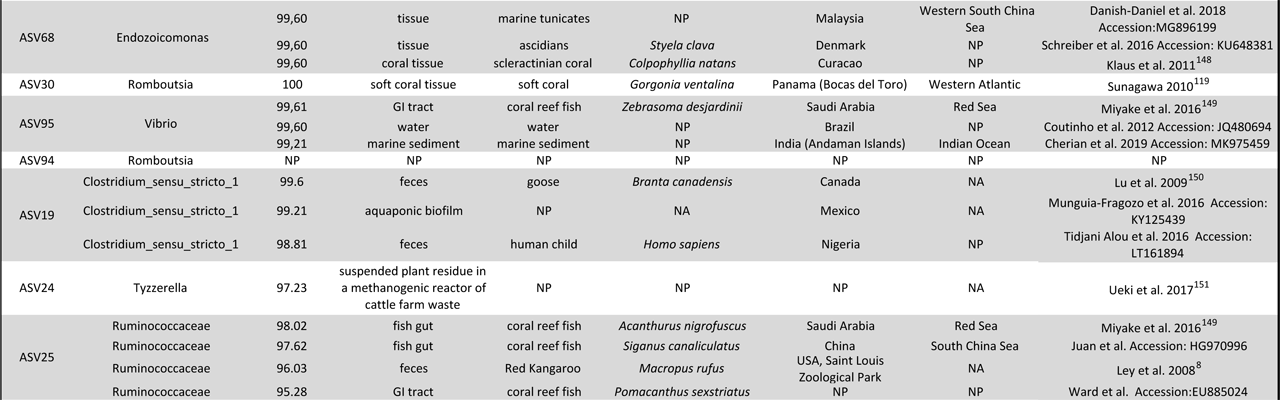
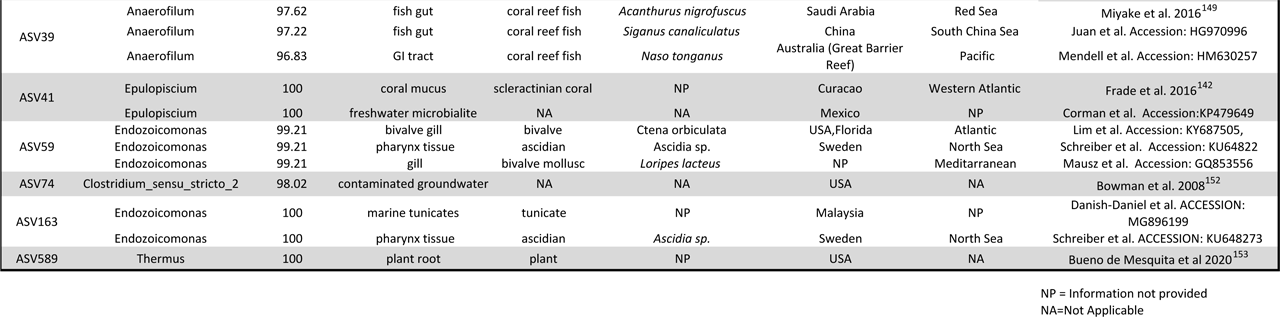
Basic Local Alignment Search Tool for nucleotides (BLASTn)^72^ search results for ASVs identified as part of the core microbiome to infer where these ASVs or close sequences have been previously identified. Core ASVs were compared to the non-redundant nucleotide (nr/nt) collection database of the National Centre for Biotechnology Information (NCBI) with BLASTn. Metadata are recorded for sequences that matched each query at 100% similarity or the first five top hits.

### Alpha diversity of the whole gut microbiome

We estimated alpha diversity using Hill numbers of three different orders of diversity (Hill numbers, {q = 0, 1, 2}) that place more or less weight on the relative abundance of ASVs. This approach allowed for balancing the representation of rare ASVs that might be the result of sequencing errors. Diversity of the gut microbiome was lower in fishes of the outer bay zone than in fishes of the inner bay and inner bay disturbed zones [Observed ASV richness (Hill number q=0); 60.23, 85.49, 75.53; Shannon index (Hill number q=1); 4.77, 7.39, 10.1; Simpson index (Hill number q=2); 2.29, 2.96, 4.58; Table S4A] (Fig. 4A, 4B and 4C). Diversity differed significantly among the three zones when taking into account ASV frequency with the Shannon index (Kruskal-Wallis-Test, *P*=0.004; Fig. 4B and Table S4A) and when emphasizing abundant ASVs with the Simpson index (Kruskal-Wallis-Test; *P*=0.013, Fig. 4C and Table S4A). However, observed ASV richness did not significantly differ among zones (Kruskal-Wallis-Test; *P* = 0.174, Table S4A) (Fig. 4A). Benjamin Hochberg corrected posthoc tests showed significantly higher Shannon diversity in fish guts of the inner versus the outer bay zone (Dunn Test; *P*=0.033, *P*=0.001). Fish of the inner bay disturbed zone had a higher microbial diversity than fishes of the outer bay zone based on both Shannon and Simpson (Dunn Test; *P*=0.004, Table S4.C). Pairwise comparisons of alpha diversity between reefs revealed that fishes resident on the reef with the highest level of coral cover overall (37.07%), Salt Creek (SCR, outer bay), had a significantly lower diversity of microbes in their guts than fishes from all three inner bay disturbed reefs (RNW, PST, PBL) for both Shannon (Dunn-Test; SCR-RNW *P*=0.013, SCR-PBL *P*=0.024) and Simpson diversity (Dunn-Test; SCR-RNW *P*=0.016, SCR-PST *P*=0.04 SCR-PBL *P*=0.026) (Table S4D).

**Figure 4.**
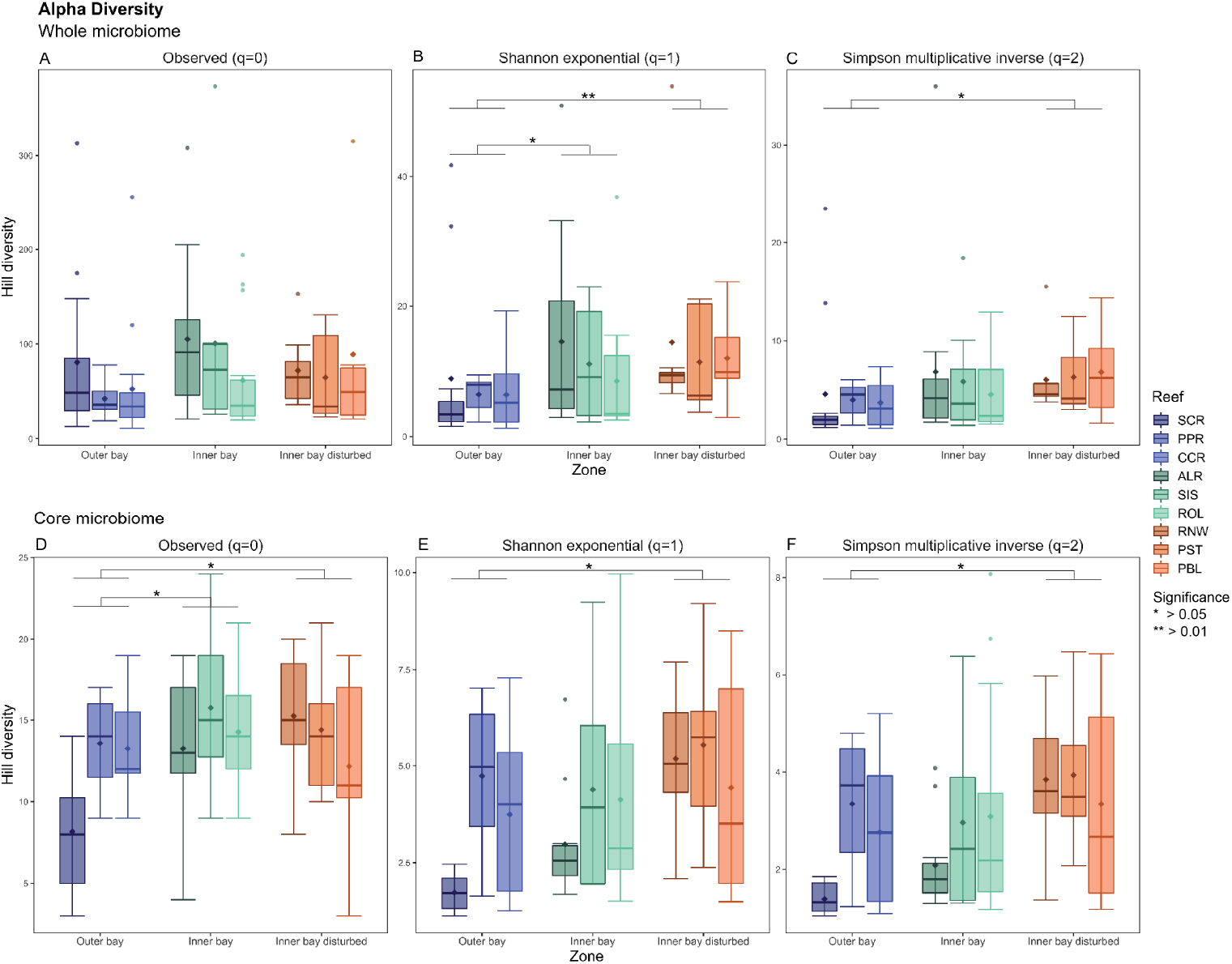
Differences in diversity of ASVs between the whole gut microbiome (A-C) and the core gut microbiome (D-F) of *Chaetodon capistratus* across reefs. Alpha diversity was measured based on Hill numbers using three metrics that put more or less weigh on common species. The observed richness (panels A and D) does not take into account relative abundances. Shannon exponential (panels B and E) weighs OTUs by their frequency. Simpson multiplicative inverse (panels C and F) overweighs abundant OTUs. Diamond shapes depict means.

### Patterns of alpha diversity of the core gut microbiome

Diversity of ASVs in the core microbiome was lowest at the outer bay when comparing ASV richness among fishes of the outer bay, inner bay, and inner bay disturbed zones [(Hill number q=0); 11.57, 14.26, 14.05] and was highest in fishes at the inner bay disturbed zone with both the Shannon index [(Hill number q=1); 2.71, 3.27, 4.45] and Simpson index [(Hill number q=2) 1.8, 2.06, 2.83] (Fig. 4D, 4E and 4F; Table S4A). Alpha diversity differed significantly among the three zones (Kruskal-Wallis-Test; observed richness: *P*=0.025; Shannon index: *P*=0.015 and Simpson index: *P*=0.016; Table S4A) and pairwise testing revealed that this was largely due to differences between fishes of the outer bay and inner bay disturbed zones (Dunn-Test with Benjamin Hochberg correction; Observed *P*=0.049; Shannon *P*=0.012; Simpson *P*=0.012, Table S4C). When comparing by reef, lower core microbial diversity in fishes from Salt Creek (SCR, outer bay) than fishes from other reefs across all zones was responsible for most significant comparisons (Table S4D).

### Beta Diversity for the whole gut microbiome

Permutational Analysis of Multivariate Dispersion (PERMDISP2) indicated no difference in variability in the whole fish gut microbiome across zones and reefs using dissimilarity metrics that put limited weight on abundant ASVs (PERMDISP2; Jaccard: *P*=0.978; modified Gower: *P*=0.182; Fig. 5A and 5B, Tables S6A and S6B). However, Bray-Curtis, which more heavily weights abundant ASVs, identified significantly higher multivariate dispersion for fishes from the inner bay disturbed zone than for fishes from the outer bay zone (PERMDISP2, *P* = 0.0007, Fig. 5C). The same pattern was observed with phylogenetic dissimilarity metrics. Only the two metrics taking into account relative abundances (i.e. GUniFrac, WUniFrac) revealed significant differences in dispersion patterns among the three zones. Using GUniFrac, an index that adjusts the weight of abundant ASVs based on tree branch lengths, gut microbiomes of fishes from the inner bay disturbed zone were significantly more spread in multivariate space than gut microbiomes of fishes from the outer bay zone (PERMDISP2, *P* = 0.021, Fig. 5E). Gut microbial communities were significantly more variable in fishes from the inner bay zone than in fishes from the outer bay zone using both GUniFrac (PERMDISP2, *P* = 0.038, Fig. 5E) and WUniFrac (PERMDISP2, *P* = 0.025, Fig. 5F).

**Figure 5.**
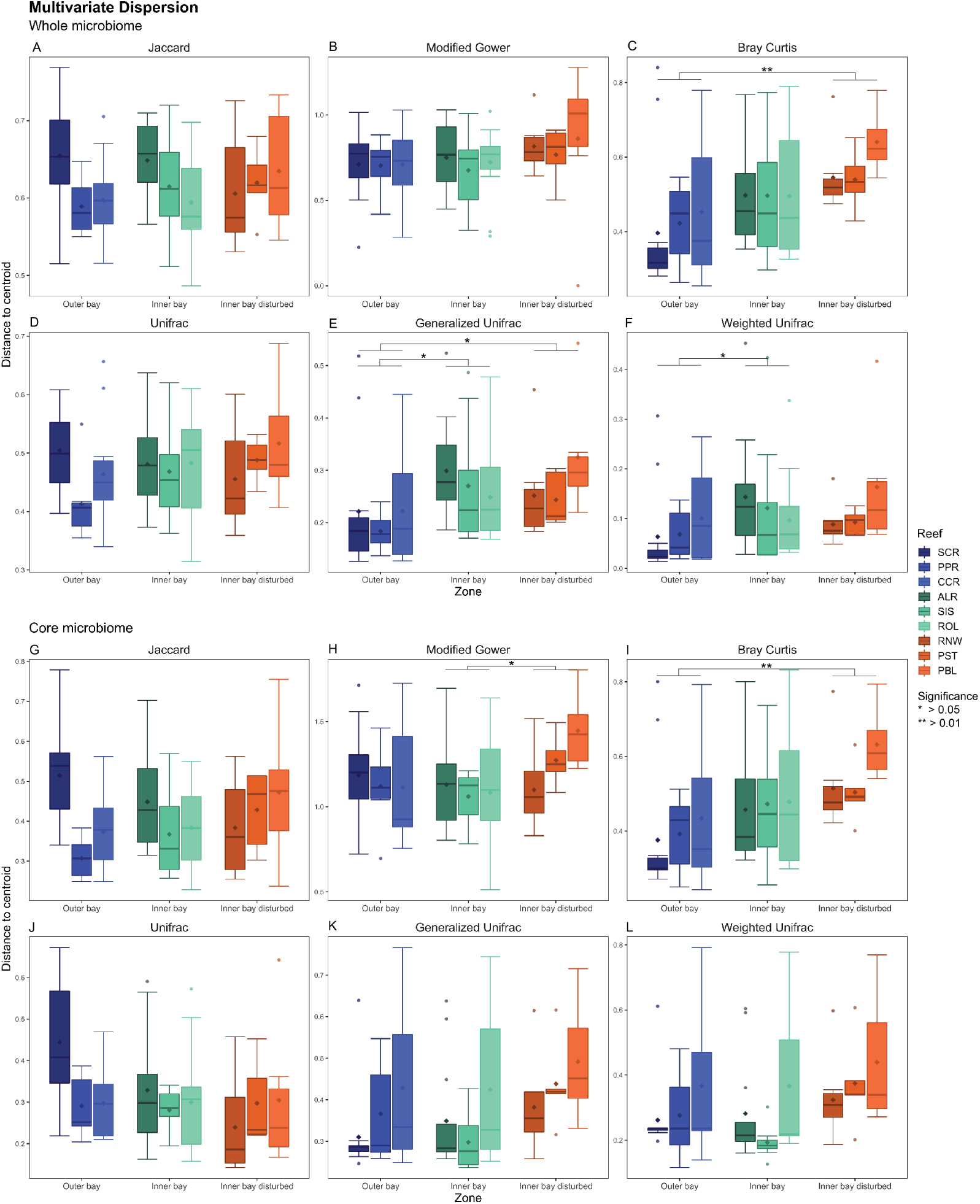
Compositional variability of the whole gut microbiome (A-F) and core gut microbiome (G-L) of *Chaetodon capistratus* across reefs. Compositional variability is measured as the distance to centroid of each group (fishes at each reef) in multivariate space. Multivariate analyses were computed with non-phylogenetic [Jaccard: panels A and G; Modified Gower: panels B and H; and Bray Curtis: panels C and I] and phylogenetic (Unifrac: panels D and J; Generalized Unifrac: E and K; Weighted Unifrac F and L) that differ in how much weigh they give to relative abundances. On one end of the spectrum, Jaccard and Unifrac only use presence-absence data, whereas on the end of the spectrum Bray Curtis and Weighted Unifrac give a lot of weigh to abundant ASVs in dissimilarity calculations. Significance depicts differences in multivariate dispersion between reef zones (ANOVA). Diamond shapes depict means.

The three Permanova models explained a small portion of the variance in the composition of the whole gut microbiome using all metrics (2.29% - 9.22%; Fig. 6A, Table S7A). Nevertheless, gut microbiome composition was significantly different between fishes from all three zones (zone model), between fishes collected inside and outside the bay (position model) and between fishes collected at inner bay reefs that differ in coral cover (cover model) when using Jaccard (Permanova; *R*^2^=0.04, *P*=0.0001; *R*^2^=0.03, *P*=0.0002; *R*^2^=0.03, *P* =0.002), modified Gower (Permanova; *R*^2^=0.06, *P*=0.0001; *R*^2^=0.04, *P*=0.0002; *R*^2^=0.04, *P*=0.001) and Bray Curtis (Permanova; *R*^2^=0.04, *P*=0.0001; *R*^2^=0.03, *P*=0.0002; *R*^2^=0.03, *P*=0.002) (Fig. 6A, Table S7A) distances. Whole gut microbiomes differed using phylogenetic metrics UniFrac (Permanova; *R*^2^=0.04, *P*=0.0004; *R*^2^=0.03, *P*=0.0007; *R*^2^=0.03, *P*=0.047) and GUniFrac (Permanova; *R*^2^=0.05, *P*=0.008; *R*^2^=0.03, *P*=0.013; *R*^2^=0.03, *P*=0.115) but not when emphasizing microbial relative read abundance (WUniFrac) (Permanova; *R*^2^=0.04, *P*=0.071; *R*^2^=0.02, *P*=0.091; *R*^2^=0.03, *P*=0.229) (Fig. 6A, Table S7A). Pairwise Adonis with Bonferroni corrected *P*-values revealed significant differences among all pairs of zones using non-phylogenetic metrics (Table S7C). Pairwise tests were significant using the Unifrac distance except between gut microbiomes of fish from the inner bay and inner bay disturbed zones. None of the pairwise tests using GUnifrac and WUnifrac were significantly different among zones (Table S7C).

**Figure 6.**
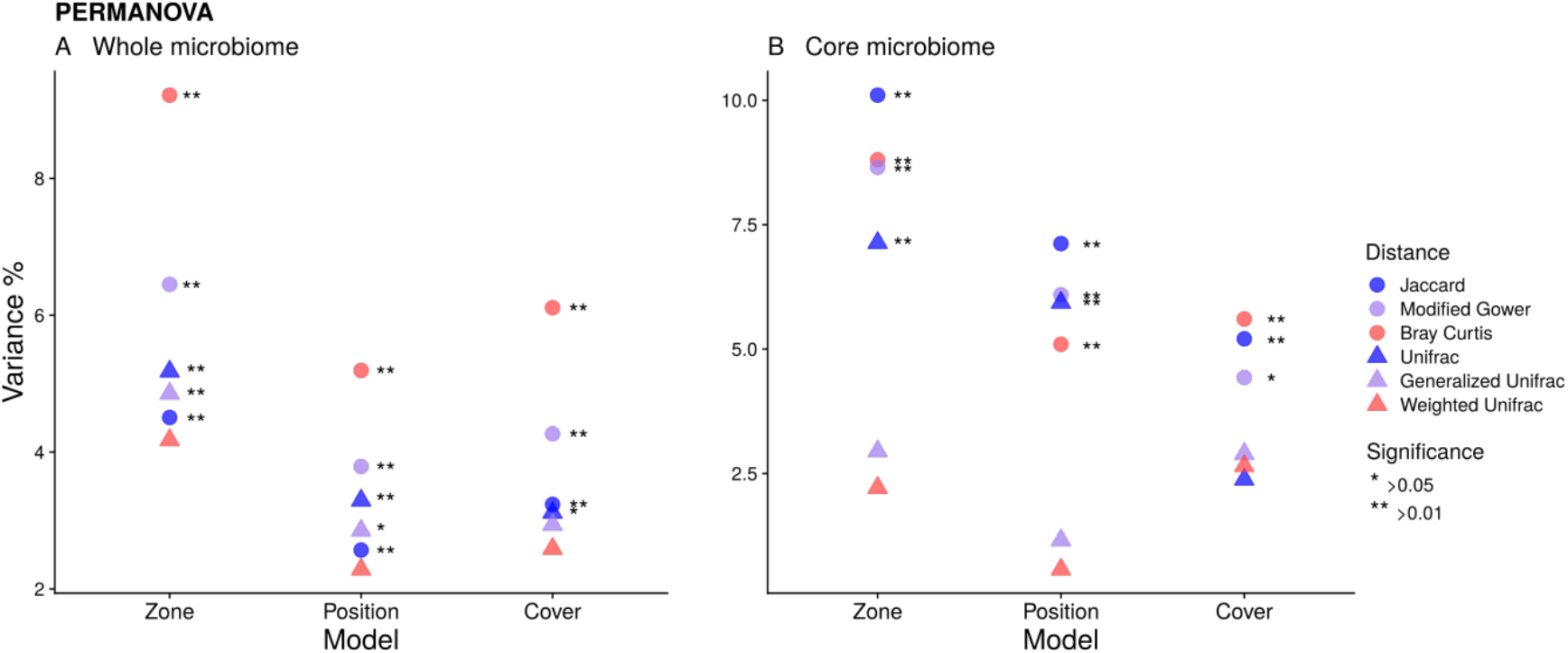
Proportion of the variance explained in Permutational Analysis of Variance (PERMANOVA) comparing the composition of the whole gut microbiome (A) and the core gut microbiome (B) of *Chaetodon capistratus*. Three independant PERMANOVA analysis were conducted. “Zone” compares gut microbiones of the three zones of the bay (inner bay, inner bay disturbed and outer bay). “Position” contrasts the composition of gut microbiones of fishes collected inside vs. outside the bay. “Cover” compares gut microbiomes of fishes on disturbed and undisturbed reefs inside the bay. Three non-phylogenetic (round shape) and three phylogenetic (triangle shape) dissimilarity metrics were used. They place more (red) or less (blue) weigh on relative abundances.

Gut microbiomes of fishes from the inner bay disturbed zone had a lower proportion of microbial reads assigned to Endozoicomonadaceae (Proteobacteria), (48.0%, 67.0%, 69.4%) but a higher proportion of Vibrionaceae (6.5%, 0.8%, 0.8%), and Rhodobacteraceae (1.0%, 0.4%, 0.4%). In contrast, the relative contribution of Spirochaetes (12.8%, 8.9%, 7.7%) and Firmicutes (20.7%, 13.5%, 16.1%) was highest in guts of fishes at the inner bay disturbed zone (Fig. S6). Within Spirochaetes, the relative abundance of Brevinemataceae was highest in gut microbiomes of fishes from the inner bay disturbed zone (13.8%, 9.1%, 7.8), while Clostrideaceae within Firmicutes contributed more to gut microbiomes of fishes at inner bay reefs but relatively little to the gut microbiomes of fishes of the outer bay zone (1.5%, 4.3%, 0.5%). Schewanellaceae (phylum Proteobacteria) represented a higher proportion of the gut microbiome of fishes at inner bay disturbed reefs (2.2%, 0.2%, 0.6%).

### Beta Diversity for the core gut microbiome

Patterns in multivariate dispersion were largely consistent between whole and core gut microbiomes. Differences among the three reef zones were significant with metrics that place more weight on ASV relative abundance (PERMDISP2; Jaccard *P*=0.83; modified Gower *P*=0.13; Bray Curtis *P*=0.005) (Fig. 5G, 5H, 5I, Table S6B). The variability of the core gut microbiome differed significantly between fishes from the inner bay and inner bay disturbed zones (PERMDISP2; modified Gower *P*=0.037) and between fishes from the inner bay disturbed and outer bay zones (PERMDISP2; Bray Curtis *P*=0.001) with highest variability levels at the inner bay disturbed zone. However, none of the phylogenetic metrics showed significant differences in dispersion among zones (PERMDISP2; Unifrac *P*=0.12; GUnifrac *P*=0.299; WUnifrac *P*=0.301) (Fig 5J, 5K, 5L, Table S6B) (Fig. 4B, Table S6B).

As with the whole gut microbiome, the three Permanova models explained a limited amount of the variance in the composition of the core gut microbiome [0.6% (position model with weighted Unifrac); 10.1% (zone model with Jaccard); Fig. 5B, Table S7B]. Yet, composition differed significantly among fish from the three zones (Permanova ‘zone model’; Jaccard *R*^2^=0.1, *P*=0.0001; modified Gower *R*^2^=0.09, *P*=0.0001; Bray Curtis *R*^2^=0.09, *P*=0.0001) and between fish at inner bay and outer bay zones (Permanova ‘position model’; Jaccard *R*^2^=0.07, *P*=0.0001; modified Gower *R*^2^=0.06, *P*=0.0001; Bray Curtis *R*^2^=0.05, *P*=0.0006) as well as between zones of differential coral cover within the bay (Permanova ‘cover model’; Jaccard *R*^2^=0.05, *P*=0.003; modified Gower *R*^2^=0.04, *P*=0.012; Bray Curtis *R*^2^=0.06, *P*=0.006) (Fig. 5B, Table S7B). The core gut microbiome appeared largely similar in composition using all phylogenetic metrics but Unifrac (Table S7B): (Permanova ‘zone model’; Unifrac *R*^2^=0.07, *P*=0.001, ‘position model’ *R*^2^=0.06, *P*=0.0001, ‘cover model’ *R*^2^=0.02, *P*=0.279). Similar to the whole microbiome, Pairwise Adonis with Bonferroni corrected *P*-values showed significant differences among almost all pairs of zones when using taxonomic metrics (Table S7D). Of the phylogenetic metrics, the only significant differences were found between the inner bay versus outer, and inner bay disturbed versus outer bay zones, with Unifrac (Table S7D). Differences in the composition of the core microbiome among reef zones was largely driven by changes in the relative abundance of ASVs assigned to the genus *Endozoicomonas* (class Gammaproteobacteria) (Fig. S7). For example, the most common *Endozoicomonas* ASV (ASV1) was much more represented in the guts of fishes of outer bay and inner bay zones than in the gut of fishes at inner bay disturbed zones (57.7%, 53.4%, 25.6%) while *Endozoicomonas* assemblages became more even towards the inner bay disturbed zone. In contrast, bacteria in the genus *Brevinema* (phylum Spirochaetes) were most abundant relative to other members of the core in fish of the inner bay disturbed zone (15.4%) and least abundant at the outer bay zones (9.6%). The giant bacterium *Epulopiscium* (family Lachnospiraceae, order Clostridia), which is known to aid the digestion of algae in surgeonfishes, contributed more to the core gut microbiome of fishes at reefs of the inner bay disturbed zone (3.5%) than the inner (1.0%) and outer bay zones (0.9%). Anaerobic, fermentative bacteria showed contrasting patterns: The relative abundance of the four Ruminococcaceae core ASVs respectively varied across reef zones (ASV15 outer 3.0%, inner 0.3%, inner disturbed 0.8%; ASV14 outer 1.9%, inner 1.2%, inner disturbed 2.2%; ASV19 outer 1.6%, inner 1.2%, inner disturbed 1.6%; ASV25 outer 0.1%, inner 1.5%, inner disturbed 1.8%), whereas *Flavonifractor* was slightly more abundant at outer reefs (outer 4.2%, inner 3.1%, inner disturbed 3.1%) (Fig. S7).

### Prevalent ASVs in each reef zone

A machine learning-based, de-noising algorithm (PIME) was used to detect sets of ASVs in the whole gut microbiome that significantly contribute to differences between reef zones. The initial out of bag (OOB) error rate (i.e., the prediction error in a RandomForest model) for our unfiltered dataset was greater than 0.1 (PIME, OOB 0.27) indicating that PIME filtering would effectively remove noise. PIME identified a prevalence cut-off of 65% for the highest improved accuracy (OOB=2.25) indicating that the model was 97.75% accurate (Table S8A). The validation step showed randomized errors (Fig. S7B) corresponded with the predicted prevalence cutoff value of 0.65 indicating absence of false positives (Type I error).

The filtered dataset after selecting ASVs that were present in at least 65% of all fish guts comprised 17 ASVs in eight families; i.e., Endozoicomonadaceae, Ruminococcaceae, Pirellulaceae, Lachnospiraceae, Brevinemataceae, Cyanobiaceae, Rhodobacteraceae, Peptostreptococcaceae (Fig. 7, Table S8A, S8B and S8C). Fish of the inner bay zone showed highest richness levels with 13 ASVs, compared to eight and nine ASVs in fish of outer bay and inner bay disturbed zones, respectively (Fig. 7). An *Endozoicomonas* ASV (ASV1), which was also a dominant component of the core, had a much higher relative abundance in fish of the outer bay zone (82.1%) than in fish of the inner bay disturbed zone (41.0%) (Fig. 7). Communities differed most in composition between fish of the outer bay and inner bay disturbed zone, whereas, fish of the inner bay zone reflected an intermediate community between these two and the highest richness of *Endozoicomonas* ASVs (N=5). Evenness among *Endozoicomonas* increased and richness decreased (3 ASVs) in fish of the inner bay disturbed zone, as observed with the core community. Bacteria in the genus *Flavonifractor* occurred in fish of both inner bay zones but not outside, whereas the outer bay zone uniquely featured Rhodobacteraceae, genus *Ruegeria*. Two distinct ASVs of the giant bacterium *Epulopiscium* (family Lachnospiraceae) were significantly prevalent in fish of the inner and inner bay disturbed zones, respectively but were more abundant at disturbed reefs (2.75%). Disturbed reefs uniquely featured anaerobic gut bacteria in the genus *Romboutsia* (family Peptostreptococcaceae) and a particular ASV in the family Lachnospiraceae (Fig. 7).

**Figure 7.**
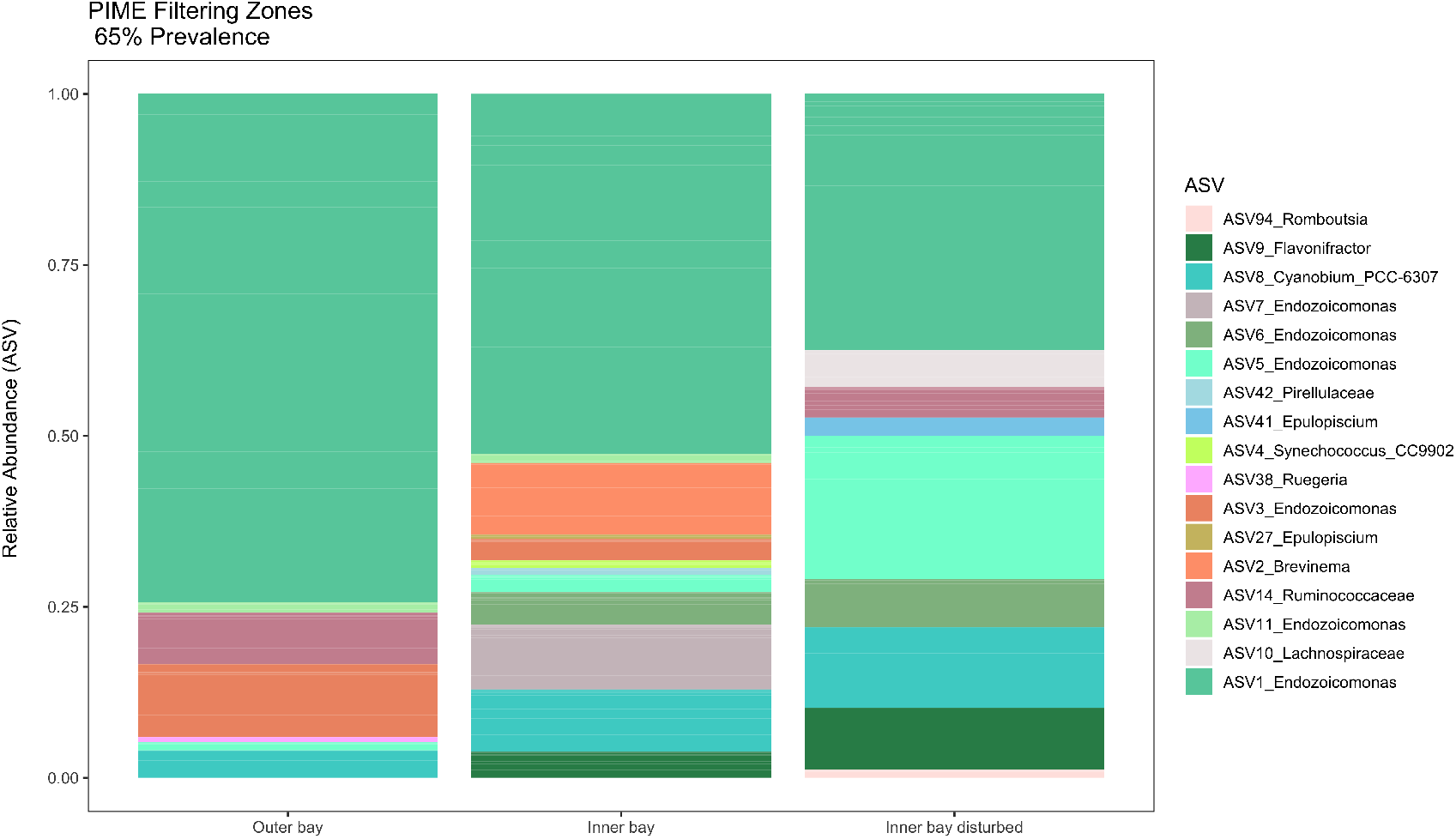
Comparison of fish gut microbiomes among three reef zones. The whole fish gut microbial dataset was filtered using Prevalence Interval for Microbiome Evaluation (PIME)^87^ to detect, which ASVs were responsible for differences among zones. Using machinelearning, PIME de-noises the data by reducing within group variability. Based on the algorithm we selected a 65% prevalence cut-off resulting in a filtered dataset of 17 ASVs at a low error rate (OOB=2.25) and high model accuracy (97.75%).

## Discussion

Detecting how host associated microbial communities differ as a function of habitat state and how spatial turnover of microbiomes varies within and among host populations is essential to understanding and predicting host species responses to environmental change. We show that both the whole and the core gut microbiome of a facultative coral feeding fish are destabilized on the most coral-depauperate reefs across a habitat degradation gradient of reefs ranging from 0% to ~30% live corals. Shifts in the fish gut microbiome may reflect changes in diet in degraded habitats and/or suggest possible limits to the host’s ability to regulate its microbiome with increasing severity of habitat degradation.

Whole gut microbial communities were significantly more diverse and variable in fish from inner bay disturbed reefs than from the outer bay zone. Conspicuously, the core microbiome, a small set of microbial strains that form sustained relationships with the fish host, also showed higher dispersion on degraded reefs, with greater variability of microbial assemblages among individual fish. Significant differences in diversity and group dispersion were only observed with diversity metrics that place less weight on rare ASVs (Fig. 4B, 4C, 5C, 5E and 5F) indicating that changes in the relative abundance of the most common taxa are responsible for this pattern. Unstable host-associated microbial communities have been observed in humans with immunodeficiency syndromes (reviewed in Williams *et al*. 2016)^34,93^ and in marine animals such as scleractinian corals and anemones under acute stress^34,94–96^. Zaneveld *et al*. (2017)^34^ referred to this pattern of variability as the “Anna Karenina principle” applied to host associated microbiomes (AKP). They argued that this is a common but often overlooked response of organisms that become unable to regulate their microbiome. Our results are consistent with patterns expected under the Anna Karenina principle suggesting that fish experience some level of stress in association with habitat degradation.

More variable gut microbial communities at disturbed reefs, where corals, the preferred food of *C. capistratus*, are nearly absent, could be a symptom of stress induced by reductions in resource availability including increased foraging costs if, for example, fish spent more energy to search, capture and handle their prey. Indeed, physiological stresses imposed by environmental conditions may cause immune signals that imbalance gut microbiota^20,34,97,98^. Disturbance to the microbiome, in turn, can affect the brain and further alter behaviours related to movement such as the ability to forage^20,99^. The scarcity of resources may also increase stress through intra- and inter-specific competition. For example, social stress in the form of aggressive interactions among conspecifics was shown to alter the behaviour and microbial assemblages associated with mice by setting off immune responses critical to host health^100^. In Indo-Pacific butterflyfishes, coral degradation was shown to decrease aggressive encounters among and within *Chaetodon* species^101^ as well as change the frequency of pair formation^102^, and the way species responded to loss of the coral resource was dependent on their level of dietary specialization^101,102^. Foraging on degraded reefs may also increase predation risk when architectural complexity is reduced^103^. Anxiety-like behaviours induced by exposure to predators can lead to sustained physiological stress in vertebrates (reviewed in Clinchy *et al*. 2013)^104^.

Another possible explanation for more variable gut microbiomes at disturbed reefs could be increased behavioural heterogeneity among fish individuals (e.g., feeding behaviour). Where preferred food sources are scarce, foraging behaviour may become more diverse and lead to increased individual specialization in various alternative food items^105,106^ translating into more varied gut microbiomes. Higher alpha diversity at the inner bay disturbed zone supports this explanation (Fig. 4B, 4C). Although *C. capistratus* is able to consume a broad range of diet items^42,107,108^, deviations from its preferred coral prey may come with fitness consequences as shown for Indo-Pacific Butterflyfishes^36,109,110^. For example, studies found that *Chaetodon* species have reduced energy reserves at reefs where they diversify or shift their diet in response to limited coral availability^36,109^. To this end, more variable gut bacterial communities at disturbed reefs in our study could be a symptom of weakened fish health due to altered nutrition.

Significant differences in the composition of the whole gut microbiome in nearly all comparisons (i.e., between all three zones, between inner and outer bay, and between inner bay disturbed and undisturbed; Fig 6A) may primarily reflect changes in diet. Microbial prevalence analysis (PIME)^87^ identified sets of ASVs that suggested a more broad, likely omnivorous trophic profile for fish where coral cover was low. This included a distinct *Endozoicomonas* community in codominance with anaerobic fermentative bacteria (e.g., *Flavonifractor* and *Romboutsia* in Firmicutes, *Epulopiscium* as well as other Lachnospiraceae in Firmicutes) (Fig. 7). Prevalence of these fermentative microbes at disturbed reefs likely reflect the consumption of algae. *Epulopiscium*, often considered a host-specific symbiont of herbivorous surgeonfishes (family: Acanthuridae)^15,18,111^, was represented in the core microbiome and identified as distinct to the inner bay predominantly at disturbed reefs. This suggests that *C. capistratus* can assimilate nutrients from algae and that this metabolic function is enhanced on degraded reefs by the increase in key microbial functional groups. Alternatively, the fish in our study may take up these microbes while foraging for invertebrates on the epilithic algal matrix. Overall, levels of *Epulopiscium* here were approximately similar to those previously found in omnivores and detritivores in the Red Sea^18^ with the two most prevalent ASVs matching (100%) to a strain previously isolated from the turf algal grazer *Naso tonganus*^112^. Additionally, the presence of Rhodobacteraceae, which are often found associated with algal biofilms^113,114^, may suggest detritus feeding but might also be related to the consumption of mucus from stressed^115^ and diseased corals^116,117^ where Rhodobacteraceae are also found. Lower relative abundance of a compositionally distinct *Endozoicomonas* community at disturbed reefs could reflect different proportions of prey species featuring *Endozoicomonas*^118^ in the diet of *C.capistratus*.

In contrast, a single dominant *Endozoicomonas* ASV along with a few Firmicutes characterized the gut microbiome of *C. capistratus* on outer bay reefs (Fig. 7). The presence of some *Endozoicomonas* ASVs shared between fish guts and potential prey (i.e., hard corals, soft corals, zoanthids, sponges) including exact matches to microbial sequences previously detected in two coral species (*Orbicella faveolata, Poritis asteroides*) at our study area in Bocas del Toro^116,119^, suggests the horizontal acquisition of these microbes via feeding on corals. In addition, we identified an ASV in the genus *Ruegeria* as indicative of outer bay reefs, which matched (100%) a sequence previously retrieved from the soft coral species *Pterogorgia anceps* on the Caribbean coast of Panama (unpublished sequence, GenBank Accession: MG099582) and which was also present across samples of potential prey taxa including hard and soft corals and sponge-infauna. Even if *Endozoicomonas* originated from the food, they might nevertheless promote the assimilation of nutrients via interactions with resident bacteria^120^.

The core microbiome composition differed under similar environmental conditions across the inner bay between fish from disturbed and undisturbed reefs (Fig. 5B). This finding suggests that bacterial communities that are most likely to have intimate metabolic interactions with *C. capistratus* might fail to provide beneficial functions to hosts at severely degraded habitats. Distinct core assemblages at the more exposed outer bay could also reflect microbial plasticity mediated by diet, gut colonization history^121^ and/or potential genetic differentiation between inner bay and outer bay fish sub-populations^122–124^.

Our analysis identified ten *Endozoicomonas* ASVs as part of the core microbiome indicating potential true resident symbionts. Members of the genus *Endozoicomonas* spp. are known as bacterial symbionts of marine sessile and some mobile invertebrates and fishes^125–128^. Reverter *et al*. (2017) found *Endozoicomonas* associated with butterflyfish gill mucus in *Chaetodon lunulatus* and Parris *et al*. (2016)^130^ found *Endozoicomonas* in the gut of damselfishes (Pomacentridae) and cardinalfishes (family: Apogonidae) pre- and to a lesser extent post-settlement on the reef. Corallivory in butterflyfishes has been found to have evolved in close association with coral reefs (Bellwood et al. 2010; Reese 1977)^35,131^ and this likely involved adaptive mechanisms to metabolize defense compounds from corals and many other sessile invertebrates (e.g., polychaetes). Adapted gut microbial communities may help butterflyfish hosts cope with toxins or facilitate the digestion of complex prey tissues as in insects, mammalian herbivores and surgeonfish^132^. It is likely that the gut microbial profile of *C. capistratus* — featuring high abundances *Endozoicomonas* — facilitates the digestion of complex coral prey. More detailed knowledge will be required to understand whether the potential intake of *Endozoicomonas* via fish browsing on sessile invertebrates is essential to trophic strategies involving fish corallivory.

Although the health of fishes is thought to be highly dependent on the state of their microbiome^10,19^, little is known about what constitutes a balanced versus imbalanced microbial assemblage. Thus, defining microbial homeostasis or dysbiosis remains challenging and these terms should be applied with caution^133–135^. We found an increase in microbiome variability, diversity, and community turnover that extended to the core microbiome suggesting that the microbiome becomes disrupted on reefs with extreme low levels of live coral cover. Additional work should focus on linking changes in the gut microbiome to host health. Our results give insight into the poorly understood spatial fluctuations in host associated microbial communities across a natural system and in response to coral reef habitat decline. This work highlights intricate links between ecosystem-scale and microbial scale processes, which have so far been mostly overlooked. We suggest there is an urgent need to integrate measurements of the role of microbes in the response of reef fishes to the global loss of coral reefs.

## Acknowledgements

We thank Lucia Rodriguez for field assistance, Joan Antaneda for her help in the laboratory, Ross Whippo for conducting the fish survey and Clare Fieseler for taking photos of the benthos. The staff of the Bocas del Toro Research Station provided logistical support. We are grateful to Kristin Saltonstall and Marta Vargas for their support at the Smithsonian Tropical Research Institute’s (STRI) Ecological and Evolutionary Genomics Laboratory. Friederike Clever was supported by a Smithsonian Short Term Fellowship. This project was funded, in part, by a grant from the Gordon and Betty Moore foundation to STRI and UC Davis (PIs: William Wcislo and Jonathan Eisen; http://doi.org/10.37807/GBMF5603) and a PhD studentship by Manchester Metropolitan University to Friederike Clever. A research permit was issued by the Ministerio de Ambiente Panamá (No. SE/A-113-17).

## Author contributions

F.C., M.L., and J.J.S. conceived the study. M.L., F.C. and R.F.P. designed the study with input from AHA. F.C., M.L., and J.J.S. conducted the fieldwork. E.C.R.G. and F.C. dissected the fish guts. F.C. extracted the DNA. J.S. and M.L. prepared the DNA for sequencing and processed the sequencing data. R.F.P., A.H.A., W.O.M. and M.L. contributed reagents and supplies. E.C.R.G. analysed the photographic benthic quadrats. F.C., J.S. and M.L. analysed the data and wrote the first draft of the manuscript with input from R.F.P. and L.G.E.W. All authors reviewed the manuscript and contributed to the final version.

