## Supplementary information for "The gut microbiome stability of a butterflyfish is disrupted on severely degraded Caribbean coral reefs"

#### I. Figures

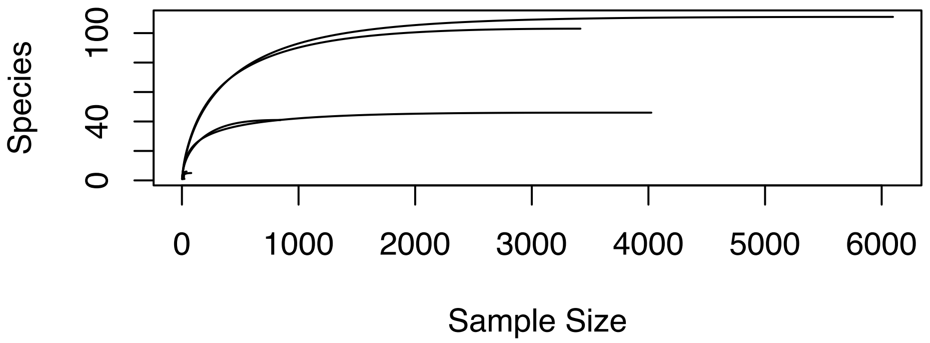

**Figure S1.** Rarefaction curves for 14 samples with fewer than 10000 sequences. Curves plateau after a few thousand reads. To limit the variability in number of sequences between samples, these samples were removed from our dataset. We rarefied the remaining samples to even sequencing depth (n = 10,369 sequences).

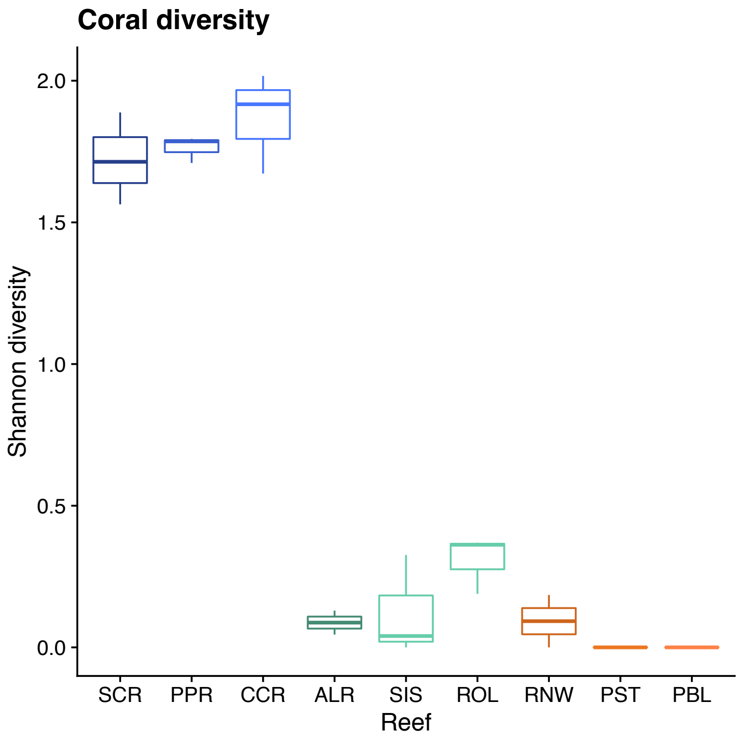

**Figure S2.** Shannon diversity of the coral community at each of nine reefs inferred from transect data. The data reflect a gradient from high coral cover at the outer bay reefs (SCR, PPR, CCR) to the inner bay (ALR, SIS, ROL) and inner bay disturbed reefs (RNW, PST, PBL).

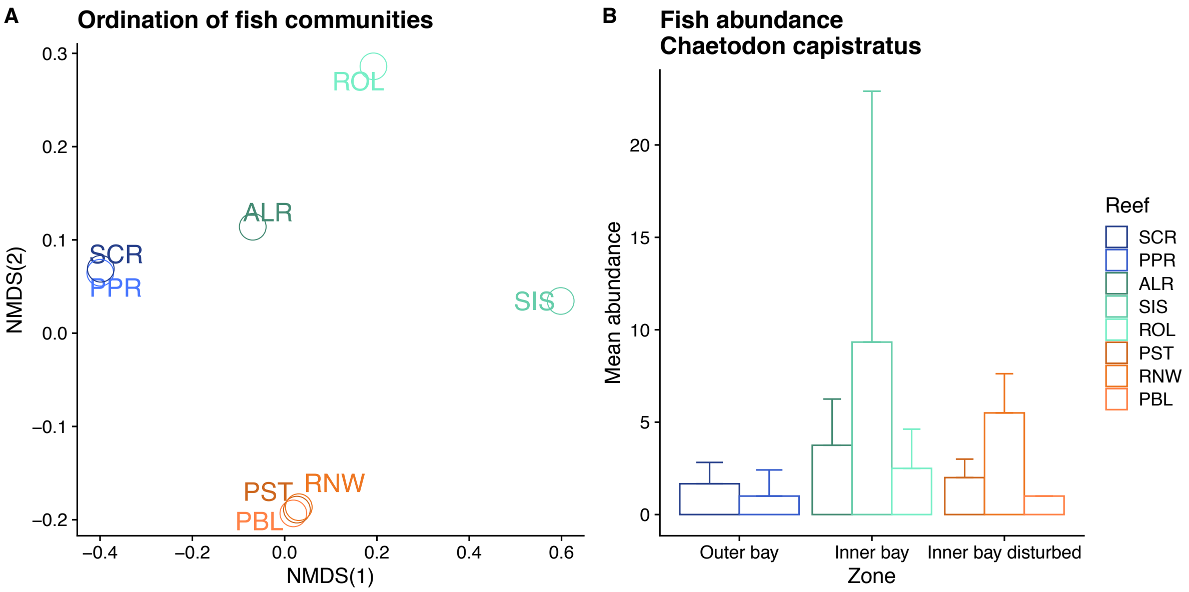

**Figure S3** (A) Differences in fish community composition among nine study reefs visualized using Nonmetric Multidimensional Scaling (NMDS) based on Bray Curtis dissimilarity. Reefs are colour-coded by reef zone: blue= outer bay, green=inner bay, orange=inner bay disturbed. (B) Study species (*Chaetodon capistratus*) abundances across nine study reefs.

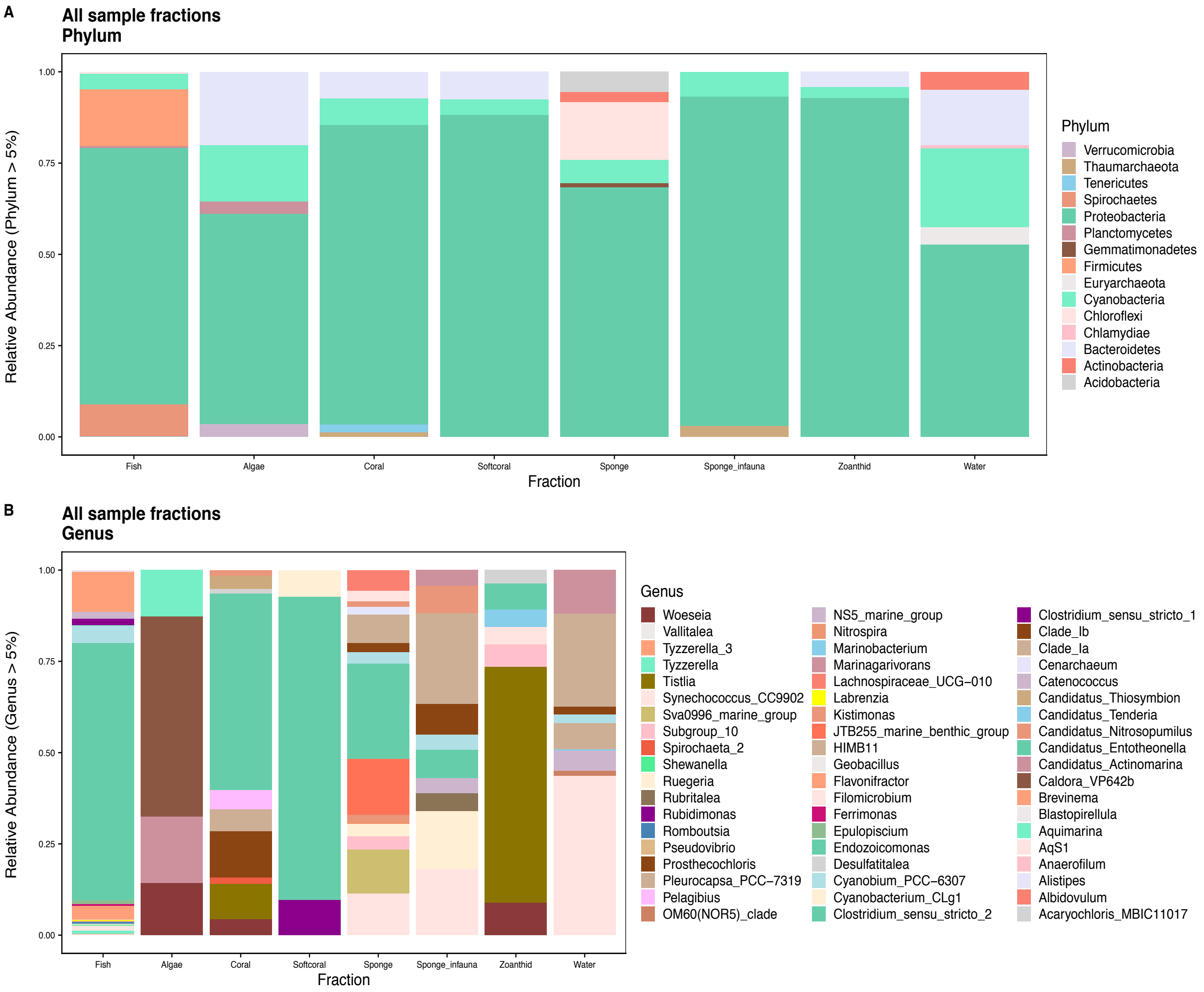

**Figure S4**. Relative abundance of microbial taxa across the different sample fractions comparing fish gut microbiome to potential prey items (i.e., algae, hard coral, soft coral, sponge, sponge infauna, zoanthid, anemone) and the surrounding seawater by phylum (A) and family (B).

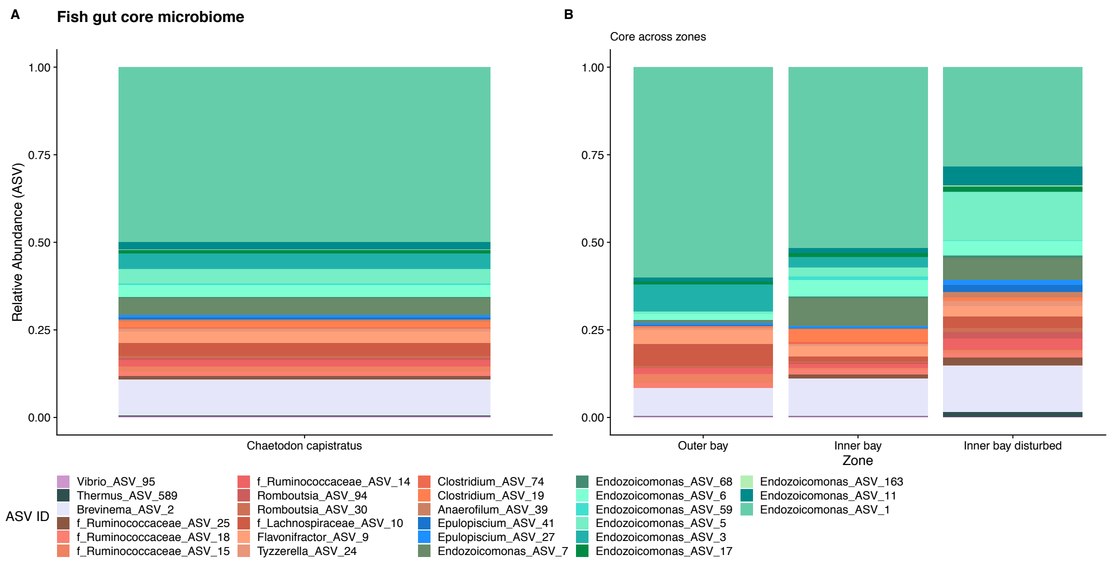

**Figure S5.** Core bacterial community in the gut of *Chaetodon capistratus* identified with Indicator Analysis by comparing 16S sequences found in fish guts to all other sample fractions (seawater and potential prey taxa) combined (A); core microbiome community variation shown across three zones (B).

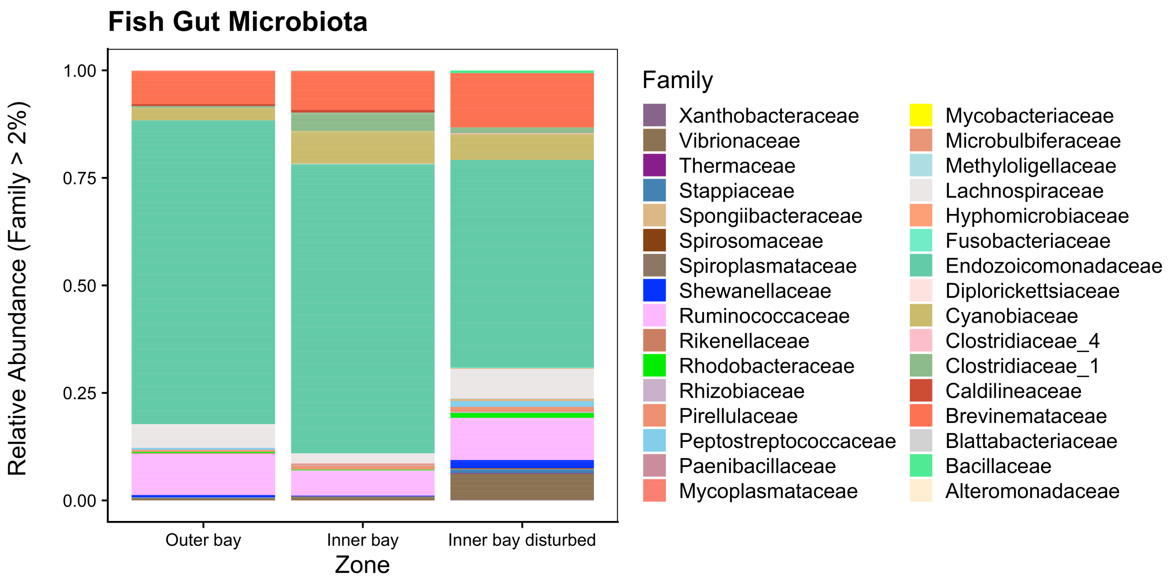

**Figure S6** Relative read abundance of bacteria in the whole fish gut microbiome across three reef zone by family

A

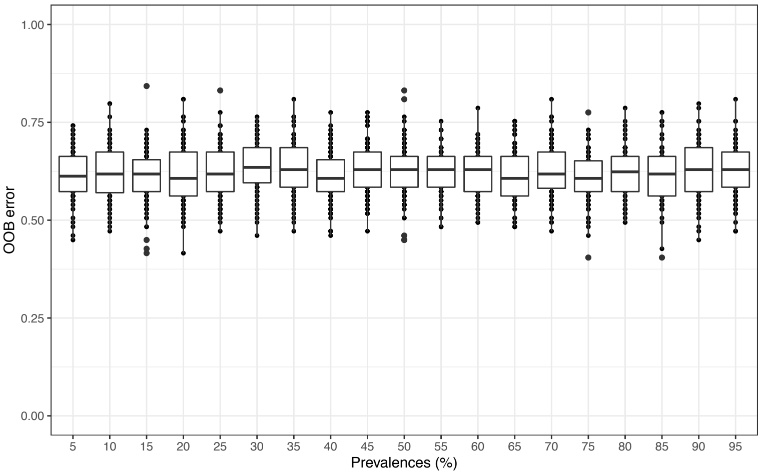

B

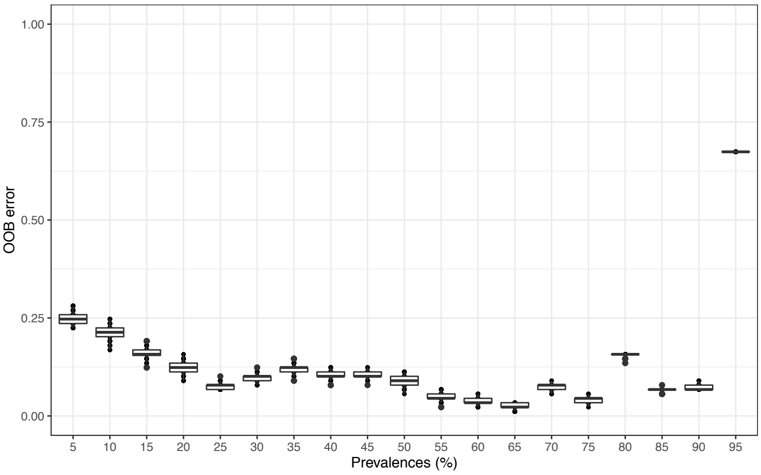

**Figure S7.** To validate PIME results, the algorithm assessed the likelihood of bias by simulating OBB error predictions. This was done by (i) by randomizing the group labels (i.e., zone identity) of the original dataset (17 ASVs) and using bootstrap aggregating (100 iterations) to perform a Monte Carlo simulation of Random Forest classifications on each filtering step (by 5 increments) generating boxplots for each prevalence interval at approximately the predicted best prevelance cut-off (65%) (A) and (ii) by repeating Random Forest OBB error estimations on the bootstrap aggregations of the filtered dataset at each prevalence interval (resulting in boxplots corresponding to the empirical data in Table 8A) (B).

### Supporting Information

#### II. Tables

**Table S1.** Number of fish individuals sampled per reef and reef site geographic coordinates. The reef zone category shows the assignment of individual reefs to zones.

| Reef | Reef name | Latitude | Longitude | Reef Zone | Number of fishes sampled |
|---|---|---|---|---|---|
| ALR | Almirante | N 09.28998 | W 82.34308 | Inner bay | 12 |
| ROL | Cayo Roldan | N 09.21478 | W 82.32454 | Inner bay | 15 |
| SIS | Cayo Hermanas | N 09.26751 | W 82.35178 | Inner bay | 8 |
| PBL | Punta Puebla | N 09.36665 | W 82.29124 | Inner bay disturbed | 6 |
| PST | Punta STRI | N 09.34885 | W 82.26292 | Inner bay disturbed | 5 |
| RNW | Runway | N 09.34193 | W 82.25997 | Inner bay disturbed | 8 |
| CCR | Cayo Corales | N 09.26747 | W 82.11983 | Outer bay | 16 |
| PPR | Popa | N 09.23344 | W 82.11189 | Outer bay | 7 |
| SCR | Salt Creek | N 09.28002 | W 82.10175 | Outer bay | 12 |

**Table S2.** Index PCR Primers

| Primer name | Illumina flow cell adapter sequence | Index sequence | Locus specific primer |
|---|---|---|---|
| SC501F | AATGATACGGCGACCACCGAGATCTACAC | ACGACGTG | ACACTCTTTCCCTACACGAC |
| SC502F | AATGATACGGCGACCACCGAGATCTACAC | ATATACAC | ACACTCTTTCCCTACACGAC |
| SC503F | AATGATACGGCGACCACCGAGATCTACAC | CGTCGCTA | ACACTCTTTCCCTACACGAC |
| SC504F | AATGATACGGCGACCACCGAGATCTACAC | CTAGAGCT | ACACTCTTTCCCTACACGAC |
| SC505F | AATGATACGGCGACCACCGAGATCTACAC | GCTCTAGT | ACACTCTTTCCCTACACGAC |
| SC506F | AATGATACGGCGACCACCGAGATCTACAC | GACACTGA | ACACTCTTTCCCTACACGAC |
| SC507F | AATGATACGGCGACCACCGAGATCTACAC | TGCGTACG | ACACTCTTTCCCTACACGAC |
| SC508F | AATGATACGGCGACCACCGAGATCTACAC | TAGTGTAG | ACACTCTTTCCCTACACGAC |
| SD501F | AATGATACGGCGACCACCGAGATCTACAC | AAGCAGCA | ACACTCTTTCCCTACACGAC |
| SD502F | AATGATACGGCGACCACCGAGATCTACAC | ACGCGTGA | ACACTCTTTCCCTACACGAC |
| SD503F | AATGATACGGCGACCACCGAGATCTACAC | CGATCTAC | ACACTCTTTCCCTACACGAC |
| SD504F | AATGATACGGCGACCACCGAGATCTACAC | TGCGTCAC | ACACTCTTTCCCTACACGAC |
| SD505F | AATGATACGGCGACCACCGAGATCTACAC | GTCTAGTG | ACACTCTTTCCCTACACGAC |
| SD506F | AATGATACGGCGACCACCGAGATCTACAC | CTAGTATG | ACACTCTTTCCCTACACGAC |
| SD507F | AATGATACGGCGACCACCGAGATCTACAC | GATAGCGT | ACACTCTTTCCCTACACGAC |
| SD508F | AATGATACGGCGACCACCGAGATCTACAC | TCTACACT | ACACTCTTTCCCTACACGAC |
| SC701R | CAAGCAGAAGACGGCATACGAGAT | ACCTACTG | GTGACTGGAGTTCAGACGTGTGCTCTTCCGATCT |
| SC702R | CAAGCAGAAGACGGCATACGAGAT | AGCGCTAT | GTGACTGGAGTTCAGACGTGTGCTCTTCCGATCT |
| SC703R | CAAGCAGAAGACGGCATACGAGAT | AGTCTAGA | GTGACTGGAGTTCAGACGTGTGCTCTTCCGATCT |
| SC704R | CAAGCAGAAGACGGCATACGAGAT | CATGAGGA | GTGACTGGAGTTCAGACGTGTGCTCTTCCGATCT |
| SC705R | CAAGCAGAAGACGGCATACGAGAT | CTAGCTCG | GTGACTGGAGTTCAGACGTGTGCTCTTCCGATCT |
| SC706R | CAAGCAGAAGACGGCATACGAGAT | CTCTAGAG | GTGACTGGAGTTCAGACGTGTGCTCTTCCGATCT |
| SC707R | CAAGCAGAAGACGGCATACGAGAT | GAGCTCAT | GTGACTGGAGTTCAGACGTGTGCTCTTCCGATCT |
| SC708R | CAAGCAGAAGACGGCATACGAGAT | GGTATGCT | GTGACTGGAGTTCAGACGTGTGCTCTTCCGATCT |
| SC709R | CAAGCAGAAGACGGCATACGAGAT | GTATGACG | GTGACTGGAGTTCAGACGTGTGCTCTTCCGATCT |
| SC710R | CAAGCAGAAGACGGCATACGAGAT | TAGACTGA | GTGACTGGAGTTCAGACGTGTGCTCTTCCGATCT |
| SC711R | CAAGCAGAAGACGGCATACGAGAT | TCACGATG | GTGACTGGAGTTCAGACGTGTGCTCTTCCGATCT |
| SC712R | CAAGCAGAAGACGGCATACGAGAT | TCGAGCTC | GTGACTGGAGTTCAGACGTGTGCTCTTCCGATCT |
| SD701R | CAAGCAGAAGACGGCATACGAGAT | ACCTAGTATGCTCTTC | GTGACTGGAGTTCAGACGTG |
| SD702R | CAAGCAGAAGACGGCATACGAGAT | ACGTACGTTGCTCTTC | GTGACTGGAGTTCAGACGTG |
| SD703R | CAAGCAGAAGACGGCATACGAGAT | ATATCGCGTGCTCTTC | GTGACTGGAGTTCAGACGTG |
| SD704R | CAAGCAGAAGACGGCATACGAGAT | CACGATAGTGCTCTTC | GTGACTGGAGTTCAGACGTG |
| SD705R | CAAGCAGAAGACGGCATACGAGAT | CGTATCGCTGCTCTTC | GTGACTGGAGTTCAGACGTG |
| SD706R | CAAGCAGAAGACGGCATACGAGAT | CTGCGACTTGCTCTTC | GTGACTGGAGTTCAGACGTG |
| SD707R | CAAGCAGAAGACGGCATACGAGAT | GCTGTAAC | GTGACTGGAGTTCAGACGTG |
| SD708R | CAAGCAGAAGACGGCATACGAGAT | GGACGTTA | GTGACTGGAGTTCAGACGTG |
| SD709R | CAAGCAGAAGACGGCATACGAGAT | GGTCGTAG | GTGACTGGAGTTCAGACGTG |
| SD710R | CAAGCAGAAGACGGCATACGAGAT | TAAGTCTC | GTGACTGGAGTTCAGACGTG |
| SD711R | CAAGCAGAAGACGGCATACGAGAT | TACACAGT | GTGACTGGAGTTCAGACGTG |
| SD712R | CAAGCAGAAGACGGCATACGAGAT | TTGACGCA | GTGACTGGAGTTCAGACGTG |

**Table S3.** Percent substrate cover across reef zones of the main substrate groups.

| | Cover % | | |
|---|---|---|---|
| **Reef Zone** | **Sponge** | **Dead hard coral** | **Live hard coral** |
| **Outer bay** | 3.1 | 17.37 | 33.46 |
| **Inner bay** | 25.24 | 10.8 | 14.65 |
| **Inner bay - disturbed** | 23.53 | 40.1 | 0.35 |

**Table S4.** Kruskal Wallis Rank Sum Test of Hill diversity[1,2]. Alpha diversity was measured using three metrics that put more or less weigh on common species (ASVs) (Hill numbers, {q = 0, 1, 2}) and Kruskal Wallis tests were used to test for significant differences in alpha diversity levels for each metric among reef zones and reefs respecitively.

**A** By Zone

| Microbiota | Factor | Diversity | Kruskal-Wallis $\chi^2$ | DF | *P*-Value |
|---|---|---|---|---|---|
| Whole | Zone | Observed q=0 | 3.494 | 2 | 0.174 |
| | | Shannon exponential q=1 | 10.996 | 2 | 0.004 |
| | | Simpsons multiplicative inverse q=2 | 8.634 | 2 | 0.013 |
| Core | Zone | Observed | 7.416 | 2 | 0.025 |
| | | Shannon exponential q=1 | 8.357 | 2 | 0.015 |
| | | Simpsons multiplicative inverse q=2 | 8.263 | 2 | 0.016 |

**B** By Reef

| Microbiota | Factor | Diversity | Kruskal-Wallis $\chi^2$ | DF | *P*-Value |
|---|---|---|---|---|---|
| Whole | Reef | Observed q=0 | 9.916 | 8 | 0.271 |
| | | Shannon exponential q=1 | 13.543 | 8 | 0.094 |
| | | Simpsons multiplicative inverse q=2 | 10.529 | 8 | 0.23 |
| Core | Reef | Observed | 19.944 | 8 | 0.011 |
| | | Shannon exponential q=1 | 23.421 | 8 | 0.003 |
| | | Simpsons multiplicative inverse q=2 | 22.052 | 8 | 0.005 |

**C**

Posthoc Dunn test by zone with Benjamin Hochberg correction

| Microbiota | Factor | Diversity | Zone | Z | *P*-Value | adjusted *P*-Value |
|---|---|---|---|---|---|---|
| Whole | Zone | Observed q=0 | Inner bay-Inner bay disturbed | 0.009 | 0.993 | 0.993 |
| | | | Inner bay-Outer bay | 1.701 | 0.089 | 0.267 |
| | | | Inner bay disturbed-Outer bay | 1.418 | 0.156 | 0.234 |
| | | Shannon exponential q=1 | Inner bay-Inner bay disturbed | -1.416 | 0.157 | 0.157 |
| | | | Inner bay-Outer bay | 2.128 | **0.033** | **0.05** |
| | | | Inner bay disturbed-Outer bay | 3.201 | **0.001** | **0.004** |
| | | Simpsons multiplicative inverse q=2 | Inner bay-Inner bay disturbed | -1.579 | 0.114 | 0.114 |
| | | | Inner bay-Outer bay | 1.587 | 0.113 | 0.169 |
| | | | Inner bay disturbed-Outer bay | 2.911 | **0.004** | **0.011** |
| Core | Zone | Observed q=0 | Inner bay-Inner bay disturbed | 0.16 | 0.436 | 0.436 |
| | | | Inner bay-Outer bay | 2.535 | **0.006** | **0.017** |
| | | | Inner bay disturbed-Outer bay | 1.966 | **0.025** | **0.049** |
| | | Shannon exponential q=1 | Inner bay-Inner bay disturbed | -1.633 | 0.102 | 0.154 |
| | | | Inner bay-Outer bay | 1.48 | 0.139 | 0.139 |
| | | | Inner bay disturbed-Outer bay | 2.875 | **0.004** | **0.012** |
| | | Simpsons multiplicative inverse q=2 | Inner bay-Inner bay disturbed | -1.945 | 0.052 | 0.078 |
| | | | Inner bay-Outer bay | 1.106 | 0.269 | 0.269 |
| | | | Inner bay disturbed-Outer bay | 2.872 | **0.004** | **0.012** |

**D**

Posthoc Dunn test by reef with Benjamin Hochberg correction (only listing significantly different comparisons)

| Microbiota | Factor | Diversity | Reef | Z | *P*-Value | adjusted *P*-Value |
|---|---|---|---|---|---|---|
| Core | Reef | Observed q=0 | SCR-SIS | -3.525 | **0.0002** | **0.008** |
| | | | SCR-ROL | 3.441 | **0.0003** | **0.01** |
| | | | SCR -PST | 2.448 | **0.007** | 0.216 |
| | | | SCR-RNW | 3.493 | **0.0002** | **0.008** |
| | | | SCR-PBL | 1.758 | **0.039** | 1 |
| | | | SCR-ALR | 2.847 | **0.002** | 0.073 |
| | | | SCR-CCR | 2.837 | **0.002** | 0.073 |
| | | | SCR-PPR | 2.483 | **0.007** | 0.202 |
| | | Shannon exponential q=1 | SCR-SIS | -2.83 | **0.005** | **0.033** |
| | | | SCR-ROL | 3.266 | **0.001** | **0.01** |
| | | | SCR-ALR | 2.252 | **0.024** | 0.11 |
| | | | SCR-RNW | 3.869 | **0.0001** | **0.004** |
| | | | SCR-PBL | 2.484 | **0.013** | 0.067 |
| | | | PST-SCR | 3.443 | **0.001** | **0.01** |
| | | | SCR-CCR | 2.794 | **0.005** | **0.031** |
| | | | SCR-PPR | 3.27 | **0.001** | **0.013** |
| | | Simpsons multiplicative inverse q=2 | ALR-PST | -1.943 | **0.052** | 0.208 |
| | | | ALR-RNW | -2.06 | **0.039** | 0.177 |
| | | | SCR-SIS | -2.417 | **0.016** | 0.081 |
| | | | SCR-ROL | 3.046 | **0.002** | **0.021** |
| | | | SCR-PBL | 2.419 | **0.016** | 0.093 |
| | | | SCR-PST | 3.385 | **0.001** | **0.013** |
| | | | SCR-RNW | 3.742 | **0.0002** | **0.007** |
| | | | SCR-CCR | 2.667 | **0.008** | 0.055 |
| | | | SCR-PPR | 3.061 | **0.002** | **0.027** |

**Table S5.** Core fish gut microbiome taxa identified with Indicator Analysis[3] comparing all fish gut microbial data combined to all other fractions combined (i.e. hard coral, soft coral, sponge, sponge infauna, zoanthid, anemone, algae, seawater). ASVs are ordered from the highest to lowest indicator value.

| ASV ID | Indicator Value | *P*-Value | Frequency | Phylum | Class | Order | Family | Genus |
|---|---|---|---|---|---|---|---|---|
| ASV.1 | 0.996349277 | 0.001 | 111 | Proteobacteria | Gammaproteobacteria | Oceanospirillales | Endozoicomonadaceae | Endozoicomonas |
| ASV.5 | 0.7413017 | 0.001 | 67 | Proteobacteria | Gammaproteobacteria | Oceanospirillales | Endozoicomonadaceae | Endozoicomonas |
| ASV.6 | 0.730337079 | 0.001 | 65 | Proteobacteria | Gammaproteobacteria | Oceanospirillales | Endozoicomonadaceae | Endozoicomonas |
| ASV.9 | 0.707730187 | 0.001 | 65 | Firmicutes | Clostridia | Clostridiales | Ruminococcaceae | Flavonifractor |
| ASV.7 | 0.696629213 | 0.001 | 62 | Proteobacteria | Gammaproteobacteria | Oceanospirillales | Endozoicomonadaceae | Endozoicomonas |
| ASV.14 | 0.696629213 | 0.001 | 62 | Firmicutes | Clostridia | Clostridiales | Ruminococcaceae | NA |
| ASV.11 | 0.674157303 | 0.001 | 60 | Proteobacteria | Gammaproteobacteria | Oceanospirillales | Endozoicomonadaceae | Endozoicomonas |
| ASV.2 | 0.659837502 | 0.001 | 71 | Spirochaetes | Spirochaetia | Brevinematales | Brevinemataceae | Brevinema |
| ASV.18 | 0.572730393 | 0.001 | 52 | Firmicutes | Clostridia | Clostridiales | Ruminococcaceae | NA |
| ASV.10 | 0.550260717 | 0.001 | 53 | Firmicutes | Clostridia | Clostridiales | Lachnospiraceae | NA |
| ASV.17 | 0.539325843 | 0.001 | 48 | Proteobacteria | Gammaproteobacteria | Oceanospirillales | Endozoicomonadaceae | Endozoicomonas |
| ASV.3 | 0.538432561 | 0.001 | 68 | Proteobacteria | Gammaproteobacteria | Oceanospirillales | Endozoicomonadaceae | Endozoicomonas |
| ASV.27 | 0.505419537 | 0.002 | 46 | Firmicutes | Clostridia | Clostridiales | Lachnospiraceae | Epulopiscium |
| ASV.68 | 0.447638938 | 0.001 | 41 | Proteobacteria | Gammaproteobacteria | Oceanospirillales | Endozoicomonadaceae | Endozoicomonas |
| ASV.15 | 0.426966292 | 0.001 | 38 | Firmicutes | Clostridia | Clostridiales | Ruminococcaceae | NA |
| ASV.30 | 0.370786517 | 0.003 | 33 | Firmicutes | Clostridia | Clostridiales | Peptostreptococcaceae | Romboutsia |
| ASV.95 | 0.348314607 | 0.002 | 31 | Proteobacteria | Gammaproteobacteria | Vibrionales | Vibrionaceae | Vibrio |
| ASV.94 | 0.34551753 | 0.004 | 32 | Firmicutes | Clostridia | Clostridiales | Peptostreptococcaceae | Romboutsia |
| ASV.25 | 0.325842697 | 0.004 | 29 | Firmicutes | Clostridia | Clostridiales | Ruminococcaceae | NA |
| ASV.19 | 0.314606742 | 0.003 | 28 | Firmicutes | Clostridia | Clostridiales | Clostridiaceae_1 | Clostridium_sensu_stricto_1 |
| ASV.24 | 0.314606742 | 0.001 | 28 | Firmicutes | Clostridia | Clostridiales | Lachnospiraceae | Tyzzerella |
| ASV.41 | 0.269662921 | 0.009 | 24 | Firmicutes | Clostridia | Clostridiales | Lachnospiraceae | Epulopiscium |
| ASV.74 | 0.258426966 | 0.008 | 23 | Firmicutes | Clostridia | Clostridiales | Clostridiaceae_1 | Clostridium_sensu_stricto_2 |
| ASV.163 | 0.251085004 | 0.005 | 24 | Proteobacteria | Gammaproteobacteria | Oceanospirillales | Endozoicomonadaceae | Endozoicomonas |
| ASV.59 | 0.235955056 | 0.006 | 21 | Proteobacteria | Gammaproteobacteria | Oceanospirillales | Endozoicomonadaceae | Endozoicomonas |
| ASV.589 | 0.235955056 | 0.006 | 21 | Deinococcus-Thermus | Deinococci | Thermales | Thermaceae | Thermus |
| ASV.39 | 0.213483146 | 0.009 | 19 | Firmicutes | Clostridia | Clostridiales | Ruminococcaceae | Anaerofilum |

**Table S6.** Multivariate beta dispersion[4,5] of fish gut microbial communities compared among reef zones. We were interested whether the variability of individual fish gut microbiomes differed among the different zones for both (A) the whole microbiome and (B) core microbiome respectively.

## A

Multivariate beta-dispersion
Whole fish gut microbiome

| Metric | Model | | Df | SumSq | MeanSq | F | N.Perm | Pr(>F) |
|---|---|---|---|---|---|---|---|---|
| Jaccard | Inner vs. Outer bay | Groups | 1 | 0.00124 | 0.0012389 | 0.34 | 10000 | 0.5637 |
| | | Residuals | 87 | 0.31702 | 0.0036439 | | | |
| | Low vs. high coral cover | Groups | 1 | 0.000012 | 0.0000116 | 0.0027 | 10000 | 0.9614 |
| | | Residuals | 52 | 0.222844 | 0.0042855 | | | |
| mod Gower | Inner vs. Outer bay | Groups | 1 | 0.051 | 0.050968 | 0.9796 | 10000 | 0.3213 |
| | | Residuals | 87 | 4.5267 | 0.052031 | | | |
| | Low vs. high coral cover | Groups | 1 | 0.11257 | 0.112573 | 2.1817 | 10000 | 0.1467 |
| | | Residuals | 52 | 2.68311 | 0.051598 | | | |
| Bray-Curtis | Inner vs. Outer bay | Groups | 1 | 0.27014 | 0.270138 | 13.215 | 10000 | 5e-04 *** |
| | | Residuals | 87 | 1.77841 | 0.020441 | | | |
| | Low vs. high coral cover | Groups | 1 | 0.09116 | 0.091155 | 2.7183 | 10000 | 0.1066 |
| | | Residuals | 52 | 1.74376 | 0.033534 | | | |
| Unifrac | Inner vs. Outer bay | Groups | 1 | 0.0058 | 0.0058003 | 0.9247 | 10000 | 0.3446 |
| | | Residuals | 87 | 0.69462 | 0.0079841 | | | |
| | Low vs. high coral cover | Groups | 1 | 0.00025 | 0.0002453 | 0.0353 | 10000 | 0.8512 |
| | | Residuals | 52 | 0.36158 | 0.0069535 | | | |
| Generalized Unifrac | Inner vs. Outer bay | Groups | 1 | 0.07265 | 0.072651 | 7.6237 | 10000 | 0.006899 ** |
| | | Residuals | 87 | 0.82908 | 0.00953 | | | |
| | Low vs. high coral cover | Groups | 1 | 0.00005 | 0.0000468 | 0.005 | 10000 | 0.9426 |
| | | Residuals | 52 | 0.48636 | 0.0093531 | | | |
| Weighted Unifrac | Inner vs. Outer bay | Groups | 1 | 0.02471 | 0.024714 | 4.8833 | 10000 | 0.0268 * |
| | | Residuals | 87 | 0.4403 | 0.0050609 | | | |
| | Low vs. high coral cover | Groups | 1 | 0.00176 | 0.0017639 | 0.2856 | 10000 | 0.6054 |
| | | Residuals | 52 | 0.32119 | 0.0061768 | | | |
| Jaccard | 3 habitat zones | Groups | 2 | 0.00017 | 0.0000857 | 0.0208 | 10000 | 0.9775 |
| | | Residuals | 86 | 0.35484 | 0.0041261 | | | |
| mod Gower | 3 habitat zones | Groups | 2 | 0.1586 | 0.07928 | 1.7449 | 10000 | 0.182 |
| | | Residuals | 86 | 3.9074 | 0.045435 | | | |
| Bray-Curtis | 3 habitat zones | Groups | 2 | 0.27062 | 0.13531 | 5.9346 | 10000 | 0.0033 ** |
| | | Residuals | 86 | 1.96081 | 0.0228 | | | |
| | Outside vs Inner disturbed | | | | | | | 0.0007 |
| Unifrac | 3 habitat zones | Groups | 2 | 0.0038 | 0.0018976 | 0.2856 | 10000 | 0.7483 |
| | | Residuals | 86 | 0.57141 | 0.0066443 | | | |
| Generalized Unifrac | 3 habitat zones | Groups | 2 | 0.06929 | 0.034647 | 3.6283 | 10000 | 0.0295 * |
| | | Residuals | 86 | 0.82122 | 0.009549 | | | |
| | outside vs inner | | | | | | | 0.0212 |
| | outside vs inner disturbed | | | | | | | 0.0381 |
| Weighted Unifrac | 3 Habitat Zones | Groups | 2 | 0.02588 | 0.0129393 | 2.561 | 10000 | **0.07729 .** |
| | | Residuals | 86 | 0.43451 | 0.0050524 | | | |
| | outside vs inner | | | | | | | 0.0251 |

Signif. codes: 0 ‘***’ 0.001 ‘**’ 0.01 ‘*’ 0.05 ‘.’ 0.1 ‘ ’ 1

**B**

Multivariate beta-dispersion
Core fish gut microbiome

| Metric | Model | | Df | SumSq | MeanSq | F | N.Perm | Pr(>F) |
|---|---|---|---|---|---|---|---|---|
| Jaccard | Inner vs. | Groups | 1 | 0.00212 | 0.0021159 | 0.1517 | 10000 | 0.7032 |
| | Outer bay | Residuals | 87 | 1.21375 | 0.0139511 | | | |
| | Low vs. high | Groups | 1 | 0.00561 | 0.0056089 | 0.3921 | 10000 | 0.5372 |
| | coral cover | Residuals | 52 | 0.74391 | 0.014306 | | | |
| mod Gower | Inner vs. | Groups | 1 | 0.0179 | 0.017936 | 0.2367 | 10000 | 0.6254 |
| | Outer bay | Residuals | 87 | 6.5933 | 0.075785 | | | |
| | Low vs. high | Groups | 1 | 0.3177 | 0.3177 | 4.5296 | 10000 | **0.0387 *** |
| | coral cover | Residuals | 52 | 3.6472 | 0.07014 | | | |
| Bray-Curtis | Inner vs. | Groups | 1 | 0.23731 | 0.237309 | 10.907 | 10000 | **0.0011 **** |
| | Outer bay | Residuals | 87 | 1.89296 | 0.021758 | | | |
| | Low vs. high | Groups | 1 | 0.07669 | 0.076693 | 3.5233 | 10000 | **0.06939 .** |
| | coral cover | Residuals | 52 | 1.1319 | 0.021767 | | | |
| Unifrac | Inner vs. | Groups | 1 | 0.0547 | 0.054697 | 3.4941 | 10000 | **0.06419 .** |
| | Outer bay | Residuals | 87 | 1.3619 | 0.015654 | | | |
| | Low vs. high | Groups | 1 | 0.01119 | 0.011193 | 0.7669 | 10000 | 0.3815 |
| | coral cover | Residuals | 52 | 0.75896 | 0.014596 | | | |
| Generalized Unifrac | Inner vs. | Groups | 1 | 0.0082 | 0.0082037 | 0.3851 | 10000 | 0.5394 |
| | Outer bay | Residuals | 87 | 1.8534 | 0.0213034 | | | |
| | Low vs. high | Groups | 1 | 0.04683 | 0.046834 | 2.3263 | 10000 | 0.1372 |
| | coral cover | Residuals | 52 | 1.04689 | 0.020132 | | | |
| Weighted Unifrac | Inner vs. | Groups | 1 | 0.00466 | 0.0046557 | 0.1563 | 10000 | 0.7019 |
| | Outer bay | Residuals | 87 | 2.59206 | 0.0297938 | | | |
| | Low vs. high | Groups | 1 | 0.07048 | 0.070476 | 2.4078 | 10000 | 0.1297 |
| | coral cover | Residuals | 52 | 1.52204 | 0.02927 | | | |
| Jaccard | 3 habitat | Groups | 2 | 0.00563 | 0.0028155 | 0.1903 | 10000 | 0.8311 |
| | zones | Residuals | 86 | 1.27258 | 0.0147974 | | | |
| mod Gower | 3 habitat | Groups | 2 | 0.3199 | 0.15996 | 2.0397 | 10000 | 0.1333 |
| | zones | Residuals | 86 | 6.7443 | 0.078422 | | | |
| | Inner disturbed vs inner | | | | | | | **0.036996** |
| Bray-Curtis | 3 habitat | Groups | 2 | 0.2554 | 0.127699 | 5.4236 | 10000 | **0.005 **** |
| | zones | Residuals | 86 | 2.0249 | 0.023545 | | | |
| | Outside vs Inner disturbed | | | | | | | **0.0014** |
| | Inner disturbed vs inner | | | | | | | 0.0675932 |
| Unifrac | 3 habitat | Groups | 2 | 0.06817 | 0.034086 | 2.1679 | 10000 | 0.1203 |
| | zones | Residuals | 86 | 1.3522 | 0.015723 | | | |
| Generalized Unifrac | 3 habitat | Groups | 2 | 0.05237 | 0.026185 | 1.2036 | 10000 | 0.2994 |
| | zones | Residuals | 86 | 1.87099 | 0.021756 | | | |
| Weighted Unifrac | 3 habitat | Groups | 2 | 0.07351 | 0.036753 | 1.2255 | 10000 | 0.3012 |
| | zones | Residuals | 86 | 2.5791 | 0.029989 | | | |

Signif. codes: 0 ‘***’ 0.001 ‘**’ 0.01 ‘*’ 0.05 ‘.’ 0.1 ‘ ’ 1

**Table S7.** Permutational Analysis of Variance (PERMANOVA) [6] results for the whole fish gut microbiome (A) and core microbiome (B). Differences among fish gut microbial communities were tested using three models: (1) among three zones (“Group” in table); (2) between reefs located inside versus outside of the bay (position model) and (3) between reefs of differential coral cover levels inside of the bay (cover model). Comparisons were done for both the whole microbiome (A) and core microbiome (B), respectively. Posthoc pairwise PERMANOVA [7] were calculated with Bonferroni corrected *P*-values for the whole (C) and core (D) communities.

## A

PERMANOVA whole fish gut microbiome

| Distance | Model | Factor | Df | SumsOfSqs | MeanSqs | F.Model | R2 | Pr(>F) |
|---|---|---|---|---|---|---|---|---|
| Jaccard | Zone/Reef | Zone | 2 | 1.561 | 0.78028 | 2.0715 | 0.04506 | **1.00E-04** |
| Jaccard | | Zone:Reef | 6 | 2.935 | 0.48922 | 1.2988 | 0.08476 | 1.00E-04 |
| Jaccard | Position/Reef | Position | 1 | 0.889 | 0.88946 | 2.361 | 0.02568 | **2.00E-04** |
| Jaccard | | Position:Reef | 7 | 3.606 | 0.51521 | 1.3678 | 0.10414 | 1.00E-04 |
| Jaccard | Cover/Reef | Cover | 1 | 0.6711 | 0.6711 | 1.7386 | 0.03235 | **0.0019** |
| Jaccard | | Cover:Reef | 4 | 2.0033 | 0.50082 | 1.3304 | 0.09657 | 0.0015 |
| modGower | Zone/Reef | Zone | 2 | 3.479 | 1.73941 | 3.0818 | 0.06451 | **1.00E-04** |
| modGower | | Zone:Reef | 6 | 5.293 | 0.88213 | 1.5629 | 0.09815 | 2.00E-04 |
| modGower | Position/Reef | Position | 1 | 2.044 | 2.0439 | 3.6213 | 0.0379 | **2.00E-04** |
| modGower | | Position:Reef | 7 | 6.728 | 0.9611 | 1.7028 | 0.12476 | 1.00E-04 |
| modGower | Cover/Reef | Cover | 1 | 1.435 | 1.43492 | 2.4181 | 0.04267 | **0.0009999** |
| modGower | | Cover:Reef | 4 | 3.707 | 0.92679 | 1.5618 | 0.1102 | 0.0016998 |
| Bray Curtis | Zone/Reef | Zone | 2 | 2.2745 | 1.13725 | 4.4843 | 0.09217 | **1.00E-04** |
| Bray Curtis | | Zone:Reef | 6 | 2.1136 | 0.35226 | 1.389 | 0.08565 | 0.049 |
| Bray Curtis | Position/Reef | Position | 1 | 1.2818 | 1.28176 | 5.0541 | 0.05194 | **0.0003** |
| Bray Curtis | | Position:Reef | 7 | 3.1063 | 0.44376 | 1.7498 | 0.12588 | 0.0029 |
| Bray Curtis | Cover/Reef | Cover | 1 | 0.9927 | 0.99273 | 3.4134 | 0.0611 | **0.0022** |
| Bray Curtis | | Cover:Reef | 4 | 1.2951 | 0.32377 | 1.1133 | 0.07971 | 0.2915 |
| Unifrac | Zone/Reef | Zone | 2 | 1.0915 | 0.54576 | 2.3968 | 0.05179 | **0.0004** |
| Unifrac | | Zone:Reef | 6 | 1.7687 | 0.29479 | 1.2946 | 0.08392 | 0.0292 |
| Unifrac | Position/Reef | Position | 1 | 0.6944 | 0.69435 | 3.0494 | 0.03294 | **0.0006999** |
| Unifrac | | Position:Reef | 7 | 2.1659 | 0.30941 | 1.3588 | 0.10276 | 0.0092991 |
| Unifrac | Cover/Reef | Cover | 1 | 0.3972 | 0.39716 | 1.69 | 0.03116 | **0.0472** |
| Unifrac | | Cover:Reef | 4 | 1.0699 | 0.26748 | 1.1382 | 0.08393 | 0.1919 |
| Generalized Unifrac | Zone/Reef | Zone | 2 | 0.3157 | 0.157865 | 2.2534 | 0.04856 | **0.008199** |
| Generalized Unifrac | | Zone:Reef | 6 | 0.5813 | 0.096875 | 1.3828 | 0.0894 | 0.055794 |
| Generalized Unifrac | Position/Reef | Position | 1 | 0.1855 | 0.185504 | 2.6479 | 0.02853 | **0.0129** |
| Generalized Unifrac | | Position:Reef | 7 | 0.7115 | 0.10164 | 1.4508 | 0.10943 | 0.0283 |
| Generalized Unifrac | Cover/Reef | Cover | 1 | 0.1302 | 0.130226 | 1.5992 | 0.0294 | 0.1154 |
| Generalized Unifrac | | Cover:Reef | 4 | 0.3903 | 0.097563 | 1.1981 | 0.08811 | 0.1997 |
| Weighted Unifrac | Zone/Reef | Zone | 2 | 0.06542 | 0.032708 | 1.9084 | 0.04175 | 0.07089 |
| Weighted Unifrac | | Zone:Reef | 6 | 0.13032 | 0.02172 | 1.2673 | 0.08318 | 0.18758 |
| Weighted Unifrac | Position/Reef | Position | 1 | 0.03581 | 0.035813 | 2.0896 | 0.02286 | 0.09119 |
| Weighted Unifrac | | Position:Reef | 7 | 0.15992 | 0.022846 | 1.333 | 0.10207 | 0.14389 |
| Weighted Unifrac | Cover/Reef | Cover | 1 | 0.0296 | 0.029603 | 1.3917 | 0.0259 | 0.229 |
| Weighted Unifrac | | Cover:Reef | 4 | 0.09229 | 0.023073 | 1.0847 | 0.08075 | 0.3582 |

## B
PERMANOVA core fish gut microbiome

| Distance | Model | Factor | Df | SumsOfSqs | MeanSqs | F.Model | R2 | Pr(>F) |
|---|---|---|---|---|---|---|---|---|
| Jaccard | Zone/Reef | Zone | 2 | 1.7548 | 0.87738 | 5.1227 | 0.10103 | **1.00E-04** |
| Jaccard | | Zone:Reef | 6 | 1.9124 | 0.31873 | 1.861 | 0.1101 | 0.0005999 |
| Jaccard | Position/Reef | Position | 1 | 1.2367 | 1.23675 | 7.2209 | 0.0712 | **1.00E-04** |
| Jaccard | | Position:Reef | 7 | 2.4304 | 0.3472 | 2.0272 | 0.13993 | 2.00E-04 |
| Jaccard | Cover/Reef | Cover | 1 | 0.518 | 0.518 | 2.9928 | 0.05207 | **0.0031** |
| Jaccard | | Cover:Reef | 4 | 1.1226 | 0.28065 | 1.6215 | 0.11284 | 0.0163 |
| modGower | Zone/Reef | Zone | 2 | 11.341 | 5.6703 | 4.4255 | 8.65E-02 | **1.00E-04** |
| modGower | | Zone:Reef | 6 | 17.216 | 2.8694 | 2.2395 | 1.31E-01 | 1.00E-04 |
| modGower | Position/Reef | Position | 1 | 7.98 | 7.9798 | 6.228 | 0.06089 | **1.00E-04** |
| modGower | | Position:Reef | 7 | 20.577 | 2.9396 | 2.2943 | 0.15701 | 1.00E-04 |
| modGower | Cover/Reef | Cover | 1 | 3.361 | 3.3608 | 2.5099 | 0.04429 | **0.0115** |
| modGower | | Cover:Reef | 4 | 8.244 | 2.0609 | 1.5391 | 0.10865 | 0.0302 |
| Bray Curtis | Zone/Reef | Zone | 2 | 1.9772 | 0.98858 | 4.2749 | 0.08802 | **1.00E-04** |
| Bray Curtis | | Zone:Reef | 6 | 1.9861 | 0.33102 | 1.4314 | 0.08841 | 0.05409 |
| Bray Curtis | Position/Reef | Position | 1 | 1.1449 | 1.14491 | 4.9509 | 0.05097 | **0.0005999** |
| Bray Curtis | | Position:Reef | 7 | 2.8183 | 0.40262 | 1.741 | 0.12546 | 0.0053995 |
| Bray Curtis | Cover/Reef | Cover | 1 | 0.8323 | 0.83225 | 3.1407 | 0.05604 | **0.005599** |
| Bray Curtis | | Cover:Reef | 4 | 1.2991 | 0.32478 | 1.2256 | 0.08748 | 0.184882 |
| Unifrac | Zone/Reef | Zone | 2 | 0.7624 | 0.3812 | 3.4988 | 0.07136 | **0.0006999** |
| Unifrac | | Zone:Reef | 6 | 1.2049 | 0.20082 | 1.8431 | 0.11278 | 0.0079992 |
| Unifrac | Position/Reef | Position | 1 | 0.6341 | 0.63405 | 5.8195 | 0.05935 | **1.00E-04** |
| Unifrac | | Position:Reef | 7 | 1.3332 | 0.19046 | 1.7481 | 0.12479 | 0.0103 |
| Unifrac | Cover/Reef | Cover | 1 | 0.1284 | 0.12835 | 1.2834 | 0.02381 | 0.2794 |
| Unifrac | | Cover:Reef | 4 | 0.462 | 0.11549 | 1.1548 | 0.08569 | 0.3073 |
| Generalized Unifrac | Zone/Reef | Zone | 2 | 0.4435 | 0.22176 | 1.3724 | 0.0295 | 0.1942 |
| Generalized Unifrac | | Zone:Reef | 6 | 1.6639 | 0.27732 | 1.7163 | 0.11068 | 0.0421 |
| Generalized Unifrac | Position/Reef | Position | 1 | 0.1747 | 0.1747 | 1.0812 | 0.01162 | 0.3097 |
| Generalized Unifrac | | Position:Reef | 7 | 1.9327 | 0.27611 | 1.7087 | 0.12856 | 0.0346 |
| Generalized Unifrac | Cover/Reef | Cover | 1 | 0.2688 | 0.26882 | 1.5667 | 0.02899 | 0.1618 |
| Generalized Unifrac | | Cover:Reef | 4 | 0.7682 | 0.19206 | 1.1193 | 0.08285 | 0.3236 |
| Weighted Unifrac | Zone/Reef | Zone | 2 | 0.2567 | 0.12835 | 1.0153 | 0.02215 | 0.38056 |
| Weighted Unifrac | | Zone:Reef | 6 | 1.2182 | 0.20304 | 1.6061 | 0.10513 | 0.09289 |
| Weighted Unifrac | Position/Reef | Position | 1 | 0.0664 | 0.066447 | 0.52562 | 0.00573 | 0.60884 |
| Weighted Unifrac | | Position:Reef | 7 | 1.4085 | 0.201211 | 1.59165 | 0.12154 | 0.08489 |
| Weighted Unifrac | Cover/Reef | Cover | 1 | 0.1903 | 0.19025 | 1.4289 | 0.0265 | 0.2238 |
| Weighted Unifrac | | Cover:Reef | 4 | 0.5972 | 0.14929 | 1.1212 | 0.08319 | 0.3352 |

**C**

Pairwise PERMANOVA

Whole fish gut microbiome

| Distance | Pairs | Df | SumsOfSqs | F.Model | R2 | p.value | p.adjusted |
|---|---|---|---|---|---|---|---|
| Jaccard | Outer vs Inner bay | 1 | 0.9179362 | 2.391052 | 0.03396813 | 0.001 | **0.003** |
| Jaccard | Outer vs Inner dist. | 1 | 0.7110085 | 1.852244 | 0.03439494 | 0.003 | **0.009** |
| Jaccard | Inner vs Inner dist. | 1 | 0.6710959 | 1.738583 | 0.03235261 | **0.002** | **0.006** |
| modGower | Outer vs Inner bay | 1 | 1.771492 | 3.2316 | 0.04536756 | **0.001** | **0.003** |
| modGower | Outer vs Inner dist. | 1 | 2.002315 | 3.31333 | 0.05990112 | **0.001** | **0.003** |
| modGower | Inner vs Inner dist. | 1 | 1.434917 | 2.3179 | 0.04267286 | **0.002** | **0.006** |
| Bray Curtis | Outer vs Inner bay | 1 | 0.7351019 | 3.061599 | 0.04308373 | **0.012** | **0.036** |
| Bray Curtis | Outer vs Inner dist. | 1 | 1.8030557 | 7.091213 | 0.12000453 | **0.001** | **0.003** |
| Bray Curtis | Inner vs Inner dist. | 1 | 0.9927345 | 3.38389 | 0.06109882 | **0.004** | **0.012** |
| Unifrac | Outer vs Inner bay | 1 | 0.7211911 | 3.130732 | 0.04401377 | **0.001** | **0.003** |
| Unifrac | Outer vs Inner dist. | 1 | 0.4669362 | 2.030862 | 0.03758708 | **0.012** | **0.036** |
| Unifrac | Inner vs Inner dist. | 1 | 0.397157 | 1.672233 | 0.03115639 | 0.056 | 0.168 |
| GUnifrac | Outer vs Inner bay | 1 | 0.1862079 | 2.693074 | 0.03809531 | **0.021** | 0.063 |
| GUnifrac | Outer vs Inner dist. | 1 | 0.1487629 | 2.294963 | 0.04226843 | **0.019** | 0.057 |
| GUnifrac | Inner vs Inner dist. | 1 | 0.1302265 | 1.575228 | 0.02940218 | 0.114 | 0.342 |
| WUnifrac | Outer vs Inner bay | 1 | 0.04234835 | 2.472368 | 0.0350828 | 0.063 | 0.189 |
| WUnifrac | Outer vs Inner dist. | 1 | 0.02331571 | 1.672923 | 0.03116885 | 0.136 | 0.408 |
| WUnifrac | Inner vs Inner dist. | 1 | 0.0296028 | 1.382646 | 0.02590066 | 0.227 | 0.681 |

**D**

Pairwise PERMANOVA

Core fish gut microbiome

| Distance | Pairs | Df | SumsOfSqs | F.Model | R2 | p.value | p.adjusted |
|---|---|---|---|---|---|---|---|
| Jaccard | Outer vs Inner bay | 1 | 1.0424578 | 5.865352 | 0.079406 | 0.001 | **0.003** |
| Jaccard | Outer vs Inner dist. | 1 | 1.0227554 | 5.475954 | 0.09527383 | 0.001 | **0.003** |
| Jaccard | Inner vs Inner dist. | 1 | 0.5180037 | 2.856284 | 0.05206849 | 0.007 | **0.021** |
| modGower | Outer vs Inner bay | 1 | 6.22975 | 4.689234 | 0.06451071 | 0.001 | **0.003** |
| modGower | Outer vs Inner dist. | 1 | 7.254523 | 4.926 | 0.0865334 | **0.001** | **0.003** |
| modGower | Inner vs Inner dist. | 1 | 3.360785 | 2.409937 | 0.04429222 | **0.019** | 0.057 |
| Bray Curtis | Outer vs Inner bay | 1 | 0.6038367 | 2.763475 | 0.03905227 | **0.012** | **0.036** |
| Bray Curtis | Outer vs Inner dist. | 1 | 1.6436521 | 7.066327 | 0.11963376 | **0.001** | **0.003** |
| Bray Curtis | Inner vs Inner dist. | 1 | 0.8322521 | 3.087106 | 0.05604045 | **0.005** | **0.015** |
| Unifrac | Outer vs Inner bay | 1 | 0.4691397 | 3.860933 | 0.05372784 | **0.004** | **0.012** |
| Unifrac | Outer vs Inner dist. | 1 | 0.5200624 | 4.280836 | 0.07606205 | **0.001** | **0.003** |
| Unifrac | Inner vs Inner dist. | 1 | 0.1283554 | 1.268345 | 0.02381048 | 0.243 | 0.729 |
| GUnifrac | Outer vs Inner bay | 1 | 0.1198879 | 0.7424537 | 0.01080051 | 0.497 | 1 |
| GUnifrac | Outer vs Inner dist. | 1 | 0.306756 | 1.734408 | 0.03227742 | 0.142 | 0.426 |
| GUnifrac | Inner vs Inner dist. | 1 | 0.2688158 | 1.5524443 | 0.02898923 | 0.164 | 0.492 |
| WUnifrac | Outer vs Inner bay | 1 | 0.05790827 | 0.4694963 | 0.00685702 | 0.664 | 1 |
| WUnifrac | Outer vs Inner dist. | 1 | 0.15775916 | 1.1256392 | 0.02118825 | 0.328 | 0.984 |
| WUnifrac | Inner vs Inner dist. | 1 | 0.1902502 | 1.4156865 | 0.0265032 | 0.213 | 0.639 |

**Table 8**. Prevalence Interval for Microbiome Evaluation (PIME)[8].

**A**

Using RandomForest, the algorithm determined the prevalence level, which provided the best model to predict differences among the three reef zone microbial communities. The table lists the output of the best prevalence function (PIME R package)[8] with the out of bag (OBB) error rate for each prevalence interval and the AVSs retained in the dataset as well as the associated number of remaining sequences (Nseq). The prevalence bin chosen for the present study (65%) is depicted in bold.

| Prevalence Interval | OBB error rate % | ASVs | Nseqs |
|---|---|---|---|
| 5% | 22.47 | 1143 | 897208 |
| 10% | 20.22 | 266 | 844546 |
| 15% | 13.48 | 118 | 799278 |
| 20% | 10.11 | 76 | 784277 |
| 25% | 8.99 | 57 | 770883 |
| 30% | 8.99 | 42 | 753340 |
| 35% | 11.24 | 36 | 744032 |
| 40% | 10.11 | 36 | 739985 |
| 45% | 8.99 | 30 | 732452 |
| 50% | 6.74 | 23 | 706945 |
| 55% | 4.49 | 21 | 674387 |
| 60% | 3.37 | 18 | 600083 |
| **65%** | **2.25** | **17** | **561593** |
| 70% | 7.87 | 10 | 496649 |
| 75% | 4.49 | 7 | 457736 |
| 80% | 15.73 | 4 | 424993 |
| 85% | 6.74 | 4 | 418457 |
| 90% | 6.74 | 3 | 405206 |
| 95% | 67.42 | 1 | 365907 |

**B**

Results for the PIME filtered community comprising 17 ASVs that are responsible for discriminating the three reef zone communities. Based on RandomForest, the algorithm calculated (i) Mean Decrease Accuracy where higher values indicate the importance of taxa in causing differences among the three reef zones, while positive values indicate ASVs truly contributed to discern zones – the importance values are reported across all decision trees and broken up by respective reef zones and (ii) Mean Decrease Gini (or mean decrease impurity), a measure of how reliable a variable predicts the model and refers to node splitting in a decision tree, where the algorithm looks for the feature (here ASV) to split where the split results in the lowest node impurity for the most optimal model.

| ASV | Inner bay | Inner bay disturbed | Outer bay | Mean Decrease Accuracy | Mean Decrease Gini |
|---|---|---|---|---|---|
| ASV6 | 0.056103703 | 0.021597619 | 0.174855597 | 0.093632368 | 6.740867519 |
| ASV7 | 0.08671229 | 0.088669841 | 0.099411424 | 0.09076883 | 7.98316477 |
| ASV3 | 0.005125934 | 0.160498485 | 0.050850058 | 0.053926743 | 5.720833825 |
| ASV9 | 0.001270734 | 0.011368326 | 0.096647221 | 0.040011572 | 3.573840343 |
| ASV42 | 0.02385976 | 0.043803535 | 0.049101124 | 0.037602644 | 4.043649685 |
| ASV27 | 0.024402986 | 0.03225267 | 0.055083597 | 0.036728284 | 3.320584212 |
| ASV41 | 0.021518957 | 0.062058802 | 0.024365059 | 0.031886283 | 3.95981884 |
| ASV2 | 0.023511911 | 0.029188745 | 0.038238022 | 0.030100204 | 3.120207217 |
| ASV10 | 0.008987344 | 0.066901371 | 0.027017528 | 0.028201424 | 3.030194539 |
| ASV4 | 0.018321354 | 0.030409235 | 0.037108942 | 0.02782255 | 2.688240385 |
| ASV14 | 0.054866006 | 0.012504834 | 0.007273479 | 0.026189242 | 2.776662362 |
| ASV38 | 0.019521715 | 0.029386797 | 0.018296985 | 0.020691546 | 2.628507801 |
| ASV94 | 0.010840017 | 0.032298196 | 0.018855767 | 0.018416502 | 2.371929934 |
| ASV1 | 0.00192246 | 0.018927056 | 0.011505233 | 0.008862312 | 1.360170402 |
| ASV11 | 0.000392347 | 0.051716667 | -0.003903 | 0.008319567 | 1.061015645 |
| ASV8 | 0.00148597 | 9.00E-05 | 0.012271321 | 0.005533648 | 1.131796017 |
| ASV5 | 3.85E-05 | 0.005655916 | 0.005317374 | 0.003572141 | 0.954048098 |

**C**

Taxonomy table of the PIME filtered community comprising 17 ASVs that are responsible for discriminating the three reef zone communities.

| ASV | Kingdom | Phylum | Class | Order | Family | Genus |
|---|---|---|---|---|---|---|
| ASV1 | Bacteria | Proteobacteria | Gammaproteobacteria | Oceanospirillales | Endozoicomonadaceae | Endozoicomonas |
| ASV2 | Bacteria | Spirochaetes | Spirochaetia | Brevinematales | Brevinemataceae | Brevinema |
| ASV3 | Bacteria | Proteobacteria | Gammaproteobacteria | Oceanospirillales | Endozoicomonadaceae | Endozoicomonas |
| ASV4 | Bacteria | Cyanobacteria | Oxyphotobacteria | Synechococcales | Cyanobiaceae | Synechococcus_CC9902 |
| ASV5 | Bacteria | Proteobacteria | Gammaproteobacteria | Oceanospirillales | Endozoicomonadaceae | Endozoicomonas |
| ASV6 | Bacteria | Proteobacteria | Gammaproteobacteria | Oceanospirillales | Endozoicomonadaceae | Endozoicomonas |
| ASV7 | Bacteria | Proteobacteria | Gammaproteobacteria | Oceanospirillales | Endozoicomonadaceae | Endozoicomonas |
| ASV8 | Bacteria | Cyanobacteria | Oxyphotobacteria | Synechococcales | Cyanobiaceae | Cyanobium_PCC-6307 |
| ASV9 | Bacteria | Firmicutes | Clostridia | Clostridiales | Ruminococcaceae | Flavonifractor |
| ASV11 | Bacteria | Proteobacteria | Gammaproteobacteria | Oceanospirillales | Endozoicomonadaceae | Endozoicomonas |
| ASV27 | Bacteria | Firmicutes | Clostridia | Clostridiales | Lachnospiraceae | Epulopiscium |
| ASV42 | Bacteria | Planctomycetes | Planctomycetacia | Pirellulales | Pirellulaceae | NA |
| ASV10 | Bacteria | Firmicutes | Clostridia | Clostridiales | Lachnospiraceae | NA |
| ASV14 | Bacteria | Firmicutes | Clostridia | Clostridiales | Ruminococcaceae | NA |
| ASV41 | Bacteria | Firmicutes | Clostridia | Clostridiales | Lachnospiraceae | Epulopiscium |
| ASV94 | Bacteria | Firmicutes | Clostridia | Clostridiales | Peptostreptococcaceae | Romboutsia |
| ASV38 | Bacteria | Proteobacteria | Alphaproteobacteria | Rhodobacterales | Rhodobacteraceae | Ruegeria |

### Supporting Information

#### III. Text

*DNA extraction of gut content*

The gastrointestinal tract of each fish was opened longitudinally to isolate the digesta and the mucosa by lightly scraping the intestinal epithelium. Between 0.05 and 0.25 g of tissue was used for DNA extraction using the Qiagen Powersoil DNA isolation kit following the manufacturers instructions with minor modifications. To improve tissue lysis, 20 µL of Proteinase K (0.4 $mg.mL^{-1}$) was added into eppendorf tubes containing power beads and solution. Samples were briefly vortexed and incubated in a shaking incubator (1000 rpm) at 60°C for 15 minutes (min). Samples were vortexed for 5 min using a Vortex Genie 2 (Scientific Industries) with vortex adapter and incubated at 60°C for 1 hour and 45 min (1000 rpm) for a total of 2 hours incubation time. DNA was eluted in 100 ul buffer (Qiagen PowerSoil kit C6 solution) and DNA concentration was quantified with a Qubit Fluorometer (dsDNA High-Sensitivity Assay Kit, Invitrogen, Life Technologies). DNA extractions of the invertebrate and macroalgal tissues (0.25g per sample) were conducted using the same modified protocol.

*PCR protocols*

The first PCR amplification was performed in a total volume of 12.5 µL with 0.2 µL of 10 millimolars (mM) forward primer 515F (5' GTGYCAGCMGCCGCGGTAA 3'; [9], 0.2 µL of 10 mM reverse primer 806R (5' GGACTACNVGGGTWTCTAAT 3'; [10]), 5 µL of "5 PRIME Hot Master Mix (2.5x)" solution, 5.1 µL of nuclease-free water and 2µL of genomic DNA extract. This combination of primers has been recommended for marine microbial studies[11]. Primers were phased with heterogeneity spacers to increase the per base variability during Illumina sequencing [12]. PCR cycling conditions were 94°C for 3 min, followed by 35 cycles at 94°C for 45 s, 50°C for 1 min and 72°C for 1 min 30 sec and a final elongation step at 72°C for 10 min. Each sample was amplified three times independently and the product checked on 1.5% agarose gel. Triplicate PCRs were then pooled and purified using paramagnetic beads (KAPA Pure Beads) at a ratio sample:beads of 1:1.6. Purified PCR products were quantified using a Qubit Fluorometer and diluted to 5 nanograms per microlitre ($ng.\mu L^{-1}$). These dilutions were used in a second PCR to add unique combinations of dual index Illumina sequencing adaptors to each sample. Each PCR amplification was performed in a total volume of 11.5 µL with 1µL of 2.5 mM Forward indexed Illumina primer, 1 µL of 2.5 mM Reverse indexed Illumina primer, 5 µL of "5 PRIMER Hot Master Mix (2.5x)" solution, 3.5 µL of nuclease-free water and 1µL of diluted DNA. Here, cycling conditions were: 94°C, 3 min, followed by 6 cycles of 94°C for 45 s, 50°C for 1 min and 72°C for 1 min 30 s; and a final elongation step at 72°C for 10 min. Unique combinations of 16 unique Forward and 24 unique Reverse Illumina indexed primers were used in order to allow multiplexing of all samples (Table S2). Finally, an equal volume of indexed PCR product for each sample was mixed into a single tube. The pool was purified two successive times with paramagnetic beads at a ratio bead:sample of 1:1 to remove leftover primers and primer dimers.
